# Optimizing C14120-based LNPs for in vitro and in vivo mRNA delivery

**DOI:** 10.1101/2025.09.02.673705

**Authors:** Ping Song, Junyi Su, Camilla Lilly Kristine Vraa, Maria Gockert, Søren Fjelstrup, Henrik Hager, Jørgen Kjems

## Abstract

Lipid nanoparticles (LNPs) have proven to be an effective delivery system for RNA therapeutics. The chemical composition of LNPs determines their functional delivery efficiency and targeting properties, which vary between in vitro and in vivo contexts. Here, we have systematically characterized and compared twenty-five novel C14120-based LNP formulations for mRNA delivery in vitro and assessed in vivo mRNA expression and biodistribution using deep sequencing of DNA barcodes in a pooled LNP-mRNA library. In vitro experiments showed correlations of lipid composition with particle size and mRNA transfection efficiency in 4 different cell lines of distinct tissue and species origin. In vivo experiments employed a pooled LNP delivery of luciferase mRNA in combination with a multiplexed barcode system, revealed strong mRNA expression after 6 hours and identified LNP compositions with organ-specific targeting properties. Individual validation of three selected LNP candidates based on mRNA expression analysis confirmed high specificity for the lung-targeting candidate, lower specificity for the liver-targeting candidate, and inconclusive results for the spleen-targeting candidate. These findings identify LNP formulations with promising potential for in vitro and in vivo organ-targeted delivery.

## Introduction

Therapeutic application of RNA is set to revolutionize the treatment of various diseases^1–4^, as recently evidenced by the success of the COVID-19 vaccine^5–9^. Two components determine the therapeutic value of any drug: the therapeutic agent and the delivery vehicle. While extensive research has focused on RNA as a therapeutic agent, including protein-coding messenger RNAs and non-coding RNAs that modulate protein levels^10,11^, equally significant efforts have been dedicated to enhancing the delivery vehicles for these therapies^12–14^.

To date, nanoparticles, particularly lipid nanoparticles (LNPs) based on ionizable lipids, dominate the realm of delivery vehicles for RNA therapeutics due to their biocompatibility and efficiency in encapsulating and delivering functional nucleic acids^15–20^. LNPs protect the RNA cargo during circulation, avoid aggregation with serum proteins, and enhance tissue targeting, extravasation, uptake, and endosomal escape^21,22^. LNPs commonly comprise ionizable lipids, cholesterol, helper lipids, and polyethylene glycol (PEG) lipids. The specific components and their ratios determine the physiochemical properties of the formulated LNP, such as surface charge, size and morphology, which in turn affect pharmacokinetics, biodistribution and tissue targeting^23,24^. Lipid formulations have been extensively studied in liver tissues, which naturally filter and uptake substances from circulation^25,26^. While recent efforts have emerged to screen LNP formulations specifically designed for delivery to the gastrointestinal and lung tissues ^27,28^, the exploration of extrahepatic mRNA delivery remains limited.

The identification of an ideal LNP formulation for a specific application, and with defined targeting properties, involves screening of numerous distinct formulations. Whereas this is feasible in vitro, it becomes time-consuming and resource-intensive in vivo ^29–32^. To address this, new innovative methods such as using DNA and mRNA barcodes for in vivo LNP library screening experiments have been developed^25,33,34^. This has enabled simultaneous evaluation of the pharmacokinetics and biodistribution of hundreds of LNPs through sequencing^33^ .

In this study, we compare in vitro and in vivo mRNA delivery efficiencies of 25 different LNPs composted of C14120 as the ionizable lipid, cholesterol, dioleoylphosphatidylcholine (DOPC) or dioleoylphosphatidylethanolamine (DOPE), and D-ɑ-tocopheryl polyethylene glycol succinate (TPGS) or 1,2-Distearoyl-sn-glycero-3-phosphorylethanolamine (DSPE)-polyethylene glycol (PEG)-2000. The commonly used helper lipids, DOPC and DOPE, have demonstrated different effects when combined with different ionizable lipids^35,36^ and facilitate distinct lipid structures within LNPs. While DOPC promotes stable laminar structures, DOPE induces fusogenic inverted hexagonal structures. The commonly used PEG lipids, TPGS and DSPE-PEG2000 (DSPE-PEG), encompass differing lengths of polyethylene glycol (PEG) tails (1000, 2000 daltons) and distinct hydrophobic head groups with vitamin E and DSPE, respectively. PEG-lipids are typically employed as a coating material and recognized for their ability to prolong half-life and reduce protein association during circulation. In addition, Pluronic F127, a triblock copolymer poly(ethylene oxide)-poly(propylene oxide)-poly(ethylene oxide) containing two PEG chains and commonly used as a surfactant, was formulated with C14120 for one LNP.

For careful evaluation of these different LNP compositions, we first characterized their properties and assessed their mRNA delivery efficiency across multiple cell lines. Subsequently, we evaluated their biodistribution in vivo using a 58-nt end-protected ssDNA barcode system and identified, as well as quantified, the specific biodistribution of LNP candidates in different mouse organs through sequencing. Finally, LNP candidates for liver, lung and spleen targeting were further evaluated individually in vivo (see Figure 1).

**Figure 1.**
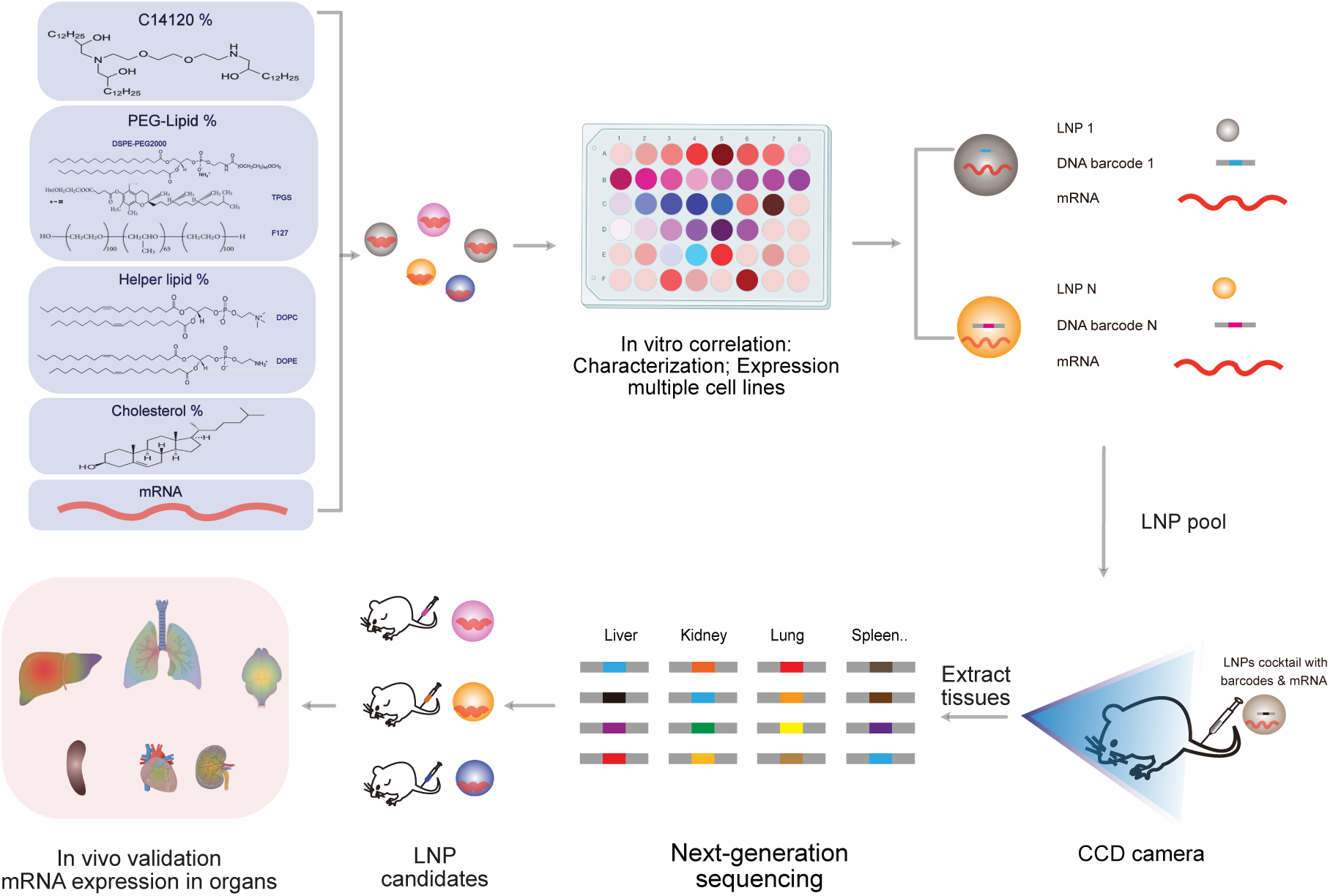
Schematic illustration of optimizing and screening of LNPs. Twenty-five types of LNPs are formulated with variations in lipid types and percentages (for details see Figure 2). After in vitro characterization and expression analysis in multiple cell lines, the LNP library is prepared by co-encapsulating mRNA and LNP-specific DNA barcodes. The libraries are then injected i.v into mice, and the biodistribution of the LNPs is assessed by deep sequencing of samples derived from the extracted tissues. Finally, a validation study is conducted by applying the individual LNP candidates to mice.

## Result

### Synthesis and characterization of C14120-based LNPs

In this study, we developed a library of 25 LNPs based on our in-house synthesized ionizable lipid, C14120^37^. C14120 was synthesized through a ring-opening reaction and purified by chromatography. The structure and molecular formula of the purified C14120 was determined by MALDI-TOF spectrum with [M+H]^+^ 787 (Figure S1a) and ^1^H NMR analysis (Figure S1b) with 3.68-3.50 (m, 11H), 3.00-2.26 (m, 10.5H), 2.23-1.90 (m, 2.6H), 1.71-0.77 (m, 75H), indicating a three-armed structure as shown in Figure S1b. We then formulated a library of LNPs (LNP1-LNP25) based on various types and percentages of C14120, helper lipid (DOPC or DOPE), cholesterol, and coated with PEG-lipids (TPGS-PEG or DSPE-PEG) as shown in Figure 2a.

**Figure 2.**
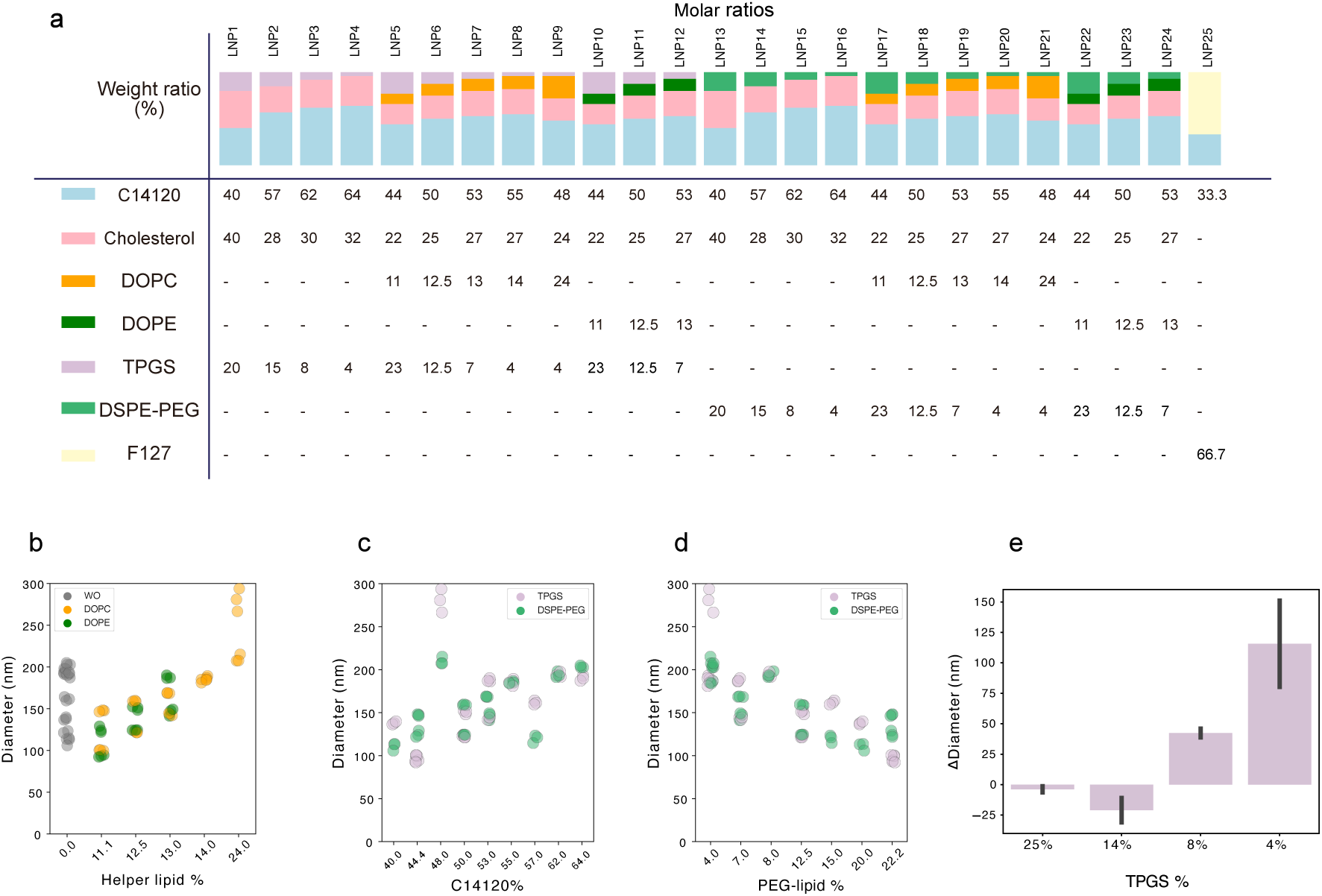
Composition and characterization of the LNPs. (a) Composition of the LNPs (weight percentages of total) (b) Influence of helper lipid (DOPE and DOPC) weight percentages on LNP size. (c) Influence of C14120 weight percentages on size of TPGS and DSPE-PEG containing LNPs. (d) Influence of PEG-lipid (TPGS and DSPE-PEG) weight percentages on LNP size. (e) Change in LNP size upon mRNA loading as a function of TPGS weight percentage (N/P ratio = 12). Data shows Mean+-SD (n = 3).

The formulated LNPs ranged in size from 100-200 nm and exhibited a positive surface charge (zeta potential) ranging from 30-70 mV (Figure S2a, b). Increased percentage of helper lipid and C14120 in the formulation generally resulted in larger particles (Figure 2b and 2c), while a rise in the percentage of PEG-lipid tended to decrease particle size (Figure 2d). Notably, LNP9 was identified as an outlier, exhibiting a larger size, likely driven by its high DOPE content in combination with TPGS as a PEG lipid. However, no significant difference was observed in the size distribution of TPGS versus DSPE-PEG coated particle library, and LNPs with and without helper lipid (Figure S2c, d). The polydispersity index (PDI) of the LNPs was generally low (<0.3), although a few LNPs exhibited values up to 0.4 (Figure S2e). The presence of any helper lipid (DOPE or DOPC) increased the PDI, albeit to a lesser degree when combined with DSPE-PEG compared to TPGS (Figure S2f).

Next, the impact of PEGylation degree on the particle size induced upon mRNA loading was assessed. We found that LNPs formulated with C14120: cholesterol: TPGS (weight ratio) ratios of 2:1:1; 4:2:1; 8:4:1; 16:8:1, exhibited a negative correlation between PEGylation levels (TPGS %) and particle size upon mRNA encapsulation. The smallest particle was observed when formulated with high percentage of TPGS (25%, 14%), while the size dramatically increased by around 140 nm with 4% TPGS (Figure 2e). The whole library of LNPs exhibited generally high encapsulation efficiency over 75% for N/P ratio of 12 (Figure S2g).

### In vitro LNP delivery across different cell types

To establish the optimal N/P ratios for mRNA expression, mRNA was loaded into three selected LNPs (LNP6, 18, and 25) through electrostatic interactions between the negatively charged mRNA and the positively charged particle surface, varying the N/P ratio between 6, 12, and 16. According to mCherry expression, N/P ratio of 12 worked best for two of the LNPs (LNP18 and 25) whereas NP ratio of 6 was more effective for LNP6 (Figure S2h). N/P ratio of 12 gave relatively high expression for all 3 LNPs and was chosen as a standard ratio for further studies (Figure S2h).

To elucidate the functional delivery of mRNA of the various LNP formulations, the individual LNPs were encapsulated with mCherry mRNA and transfected into human-derived cell lines Hep G2, HEK293-H and ARPE-19, and mouse-derived cell lines AML12 and bEnd.3 (Figure S3a-f).

First, we compared the influence of different types of PEG-lipid (TPGS and DSPE-PEG) as coating material on mRNA expression efficiency. bEnd.3 and AML12 cells were transfected with formulation ratios of C14120:cholesterol:DOPC:PEG-lipid (either TPGS or DSPE-PEG) of 4:2:1:1 (LNP6 and LNP18) (Figure 3a-c), and lipofectamine 2000 (LIPO) as a positive control. TPGS-based formulation showed higher expression in AML12 compared to DSPE-PEG-based formulation, while DSPE-PEG based formulation showed elevated expression in mouse endothelial bEnd.3 cell line. This suggests distinct cellular preferences for surface molecule compositions, which were further investigated using the full LNP library repertoire below.

**Figure 3.**
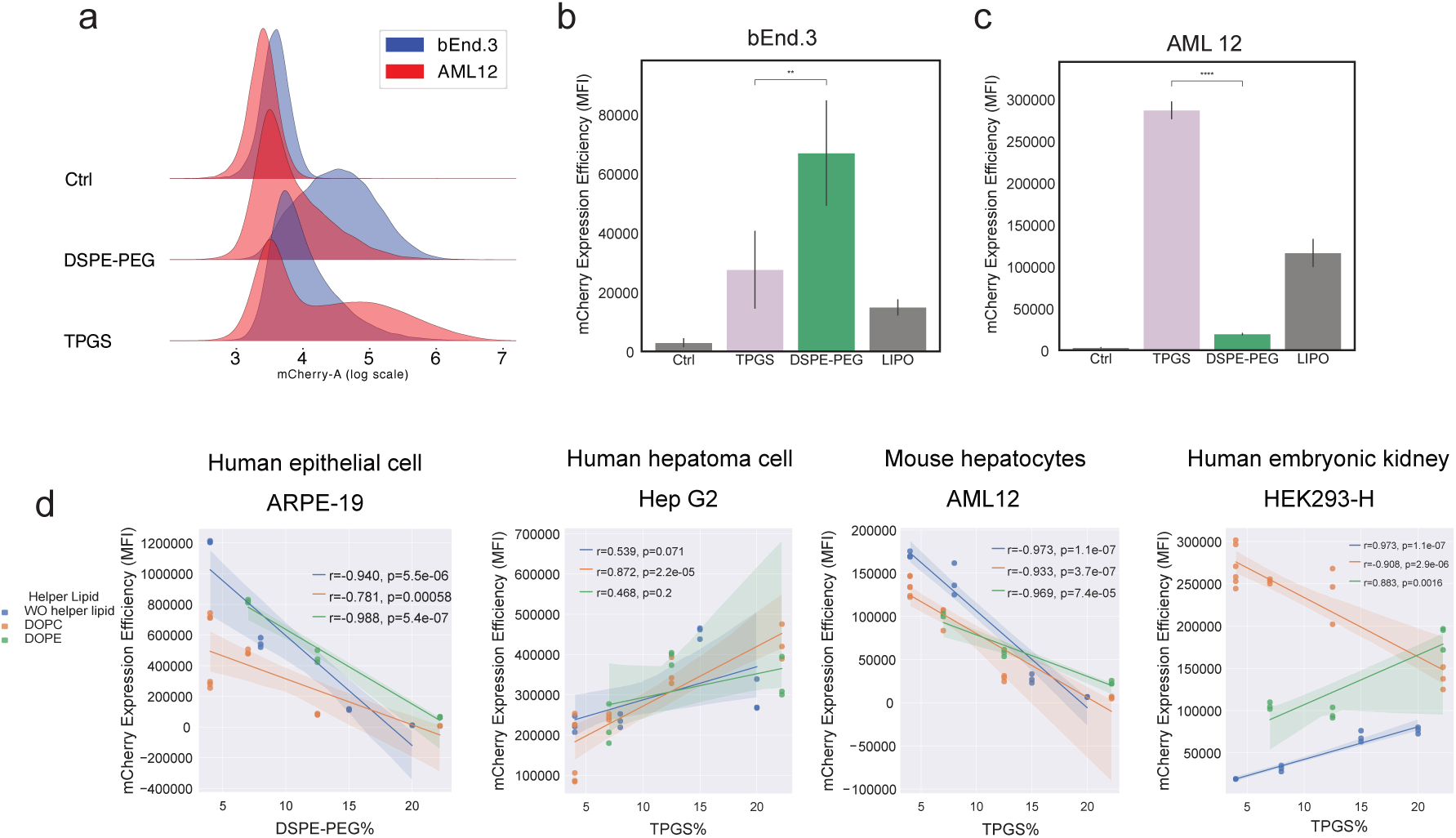
Cell-type and origin-dependency of in vitro mCherry expression for individual LNPs measured 24 hours post-transfection. (a) Histograms showing mCherry expression in bEnd3 (blue) and AML12 (red) cells upon transfection with LNPs formulated with different types of PEG-lipids (TPGS and DSPE-PEG) in comparison to respective untransfected sample (Ctrl). (b-c) mCherry expression levels, respective of data shown in a), but including a Lipofectamine 2000 (LIPO) transfection control. (c) mCherry expression level as a function of PEG-lipid percentages (DSPE-PEG and TPGS) in ARPE-19, Hep G2, AML12 and HEK293-H. Data are presented as Mean+-SD (n = 3); * *p < 0.05*, ** *p < 0.01*, *** *p < 0.001*. Significance was determined by independent *t*-test.

The TPGS-coated LNPs 1-12 were transfected into Hep G2, AML12, and HEK293-H and the DSPE-PEG coated LNPs 13-24 were transfected into ARPE-19 cells and bEnd.3 cells (Figure S3a-f), allowing for comparative analysis of similar cell types with different species using PEG-lipid-coated LNP libraries tailored for the cellular preferences shown above. The different cell lines exhibited distinct expression correlations to the increase in PEG-lipids dependent on their species and tissue origin (Figure 3d). Human-derived cell lines ARPE-19 and Hep G2 displayed inverse correlations which may be due to the different tissue origin or difference in PEG-lipid type in the formulations, while liver cells Hep2G and AML12 exhibited species-specific variations (Figure 3d). Notably, in HEK293-H cells, the expression correlation with increased PEG percentage varies with different helper lipid types. Alongside the PEG-lipid percentage, the levels of C14120 exhibited a positive correlation with mRNA expression in ARPE-19 and ALM12 cells, whereas no statistically significant correlation was found in Hep G2 cells (Figure S3g). Furthermore, the correlation between mRNA expression and types and amount of helper lipid types was not evident, primarily due to the limited helper lipid levels in the formulations (Figure S3h).

To elucidate the correlation between mRNA expression and cellular uptake, HEK293-H and AML12 cells were co-transfected with mRNA encoding EYFP and a DNA barcode (Barcode-ATTO647), each encapsulated in individual LNPs from LNP1-24, and compared with SM102. The codelivery results correlated with the previously observed trend in mRNA expression, indicating that the incorporation of DNA barcode did not exhibit any significant impact on the transfection profile and EYFP expression of the LNPs. In HEK293-H cells with typical high transfection efficiency^38^ , demonstrated lower cellular uptake, while exhibited significantly higher mRNA expression than AML12 cells. A comparison of mRNA expression levels and cellular uptake reveals that while cellular uptake is essential for expression, it does not necessarily correlate with mRNA expression levels, particularly across different LNP formulations and cell lines. Moreover, certain LNP formulations demonstrated higher expression levels compared to SM102 in HEK293-H cells. In contrast, in the more challenging-to-transfect AML12 cell line, the majority of LNP formulations exhibited superior expression and uptake levels relative to SM102 (Figure S4a-d).

Generally, LNP formulations showed cell-specific mRNA expression, with significant influence by the lipid types and levels. Notably, mRNA expression levels were not necessarily aligned with cellular uptake levels. In particular, the PEG-lipid demonstrated distinct cell specific preferences (summarized in Table 1).

**Table 1.** Summary of effective LNPs for mRNA expression in various cell lines and the cell preference for PEG-lipid types.

| Cell line | LNP number | Preference for PEG-lipid types |
| --- | --- | --- |
| Hep G2 | 2, 5, 11 | Not tested |
| AML12 | 4, 8, 12 | TPGS |
| ARPE-19 | 16, 20, 24 | TPGS |
| HEK293-H | 6, 7, 8, 9 | Not tested |
| bEnd.3 | 16, 19, 24 | DSPE-PEG |

### In vivo performance of LNP library

To simultaneously evaluate the biodistribution and mRNA expression in vivo, different barcode-containing ssDNA were added to each LNP before mixing them into a pool. Each barcode sequence contained 4 random nts to monitor PCR bias, followed by a unique 8nt-barcode sequence providing the LNP ID. The 8nt-barcode sequence was provided in Table S1.^33^ Two terminal primer binding sites were included to enable PCR amplification for quantification in the tissue (Figure 4a). End protection and internal phosphorothioate modification protected the barcode against enzymatic degradation.

**Figure 4.**
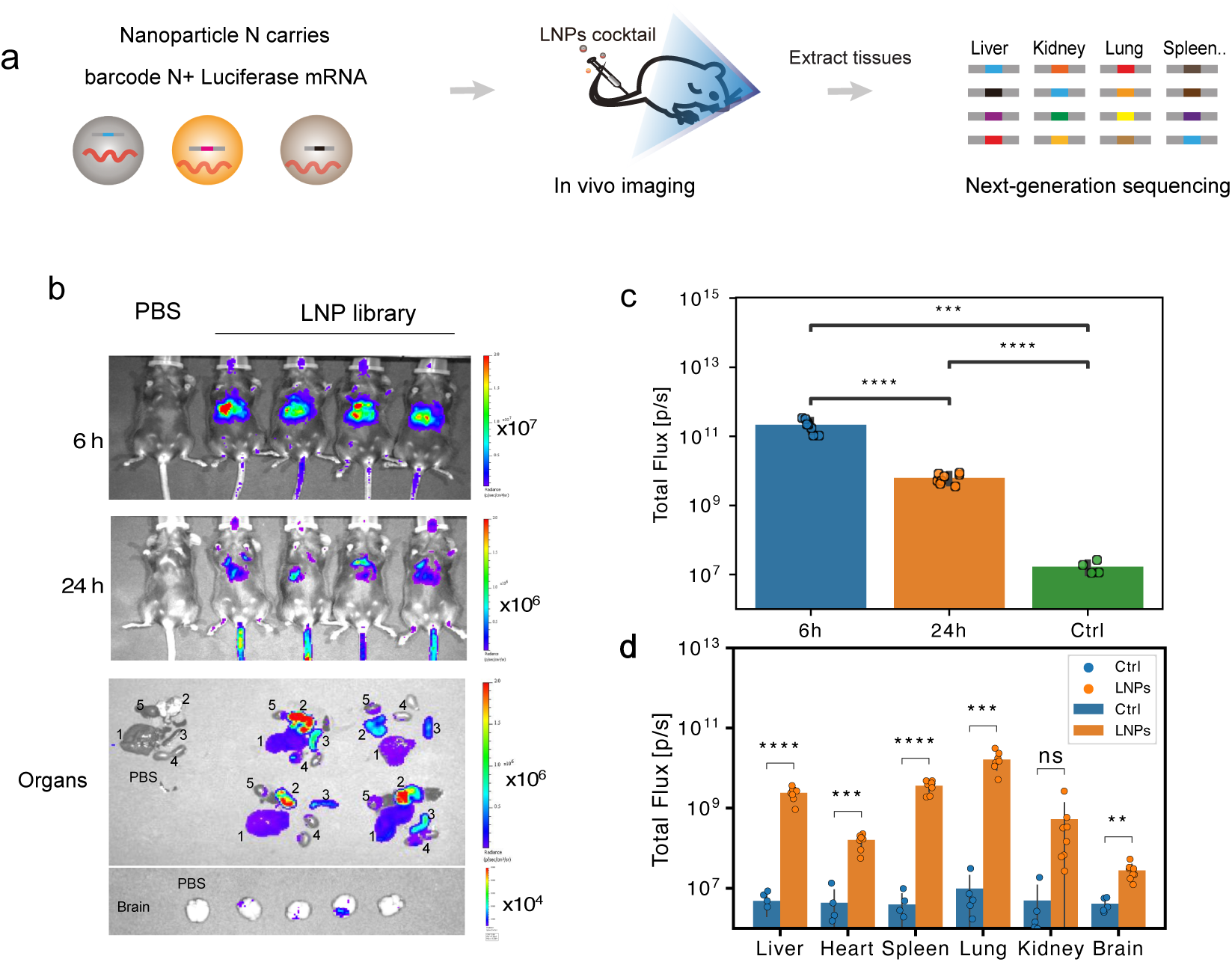
In vivo delivery of the barcoded LNP library encapsulated with Luc-mRNA enabled the detection of both mRNA expression from the library of LNPs and the identification of specific LNPs by subsequent barcode sequencing from individual organs. (a) Illustration of intravenous injection of the barcoded LNP library co-encapsulated with Luc-mRNA and DNA barcodes in mice, that subsequently allows tracking of individual LNPs by tissue-specific sequencing (b) Representative Luc-mRNA expression in live mice (6 h and 24 h) and organs (only 24 h) imaged by IVIS after i.p. injection of D-luciferin. Organ types were denoted: 1: Liver; 2: Lung; 3: Spleen; 4: Kidney; 5: Heart. Brain samples are shown below at 100 times higher sensitivity threshold. (c) Quantification of the total flux signal in live mice and (d) organs. Data are presented as Mean+-SD (LNP group n = 8, PBS group n = 5); * *p < 0.05*, ** *p < 0.01*, *** *p < 0.001*. Significance was determined by independent *t*-test.

To exclude content mixing by LNP fusion when mixed into a library, a FRET-based assay was conducted between Cy3-siRNA loaded LNPs (LNP/Cy3-siRNA) and Cy5-siRNA loaded LNPs (LNP/Cy5-siRNA). LNP19 and LNP25 were selected as representative particles and encapsulated with either Cy3-siRNA, Cy5-siRNA or a combination of both. Upon mixing, no obvious variations of Cy5/Cy3 ratio compared to free siRNAs was detected. In contrast, co-encapsulation of Cy3-siRNA and Cy5-siRNA showed significantly higher Cy5/Cy3 ratios indicative of the FRET signal (Figure S5a). We therefore conclude that the content mixing between LNPs in the mixed library is neglectable.

Mice (n=8) were intravenously injected with the barcoded LNP library and expression of Luc-mRNA was monitored at 6 and 24 hours. A pronounced bioluminescence signal was observed, predominantly in the liver after 6 hours (Figure 4b), maintaining detectable signals with statistical significance in the total flux across the entire body at 24 hours (Figure 4c). Notably, at 24 hours, the signal intensity in lung tissue was stronger than the liver signal. The bioluminescence from isolated liver, spleen, lung, heart and brain tissue exhibited significantly higher signals compared to control mice after 24 hours (n = 5; Figure 4d).

To evaluate the biodistribution of LNPs, nucleic acids were isolated from the lung, liver, spleen, heart, kidney and the brain, followed by PCR-amplification of the LNP-specific barcodes with tissue-specific primers (Table S2) and next-generation sequencing. Based on the distribution barcodes, we observed distinct and reproducible LNP signatures in the different organ samples (Figure 5a and b). Several LNPs exhibited similar organ preferences, while some LNPs appeared to preferentially target specific tissues. Importantly, due to the deviations in barcode purification and higher PCR amplification cycles for samples with low accumulation (heart and brain, 5 cycles more), no direct comparison across the different organs is possible (Figure S6a-f). Some LNPs were more highly represented in multiple organs, such as LNP11 (although predominantly in liver) and LNP13, whereas other LNPs were enriched in individual organs, such as LNP8 in the lung, LNP5 in the brain and LNP14 in the spleen. Finally, some LNPs, such as LNP12 or LNP20, exhibited generally low representation in multiple organs. Notably, while some distinct profiles are observed, highly represented LNPs, particularly LNP11, hamper the identification of general targeting trends concerning lipid types and percentages (Figure S7). A principal component analysis (PCA) visualized a distinct separation of the patterns of applied LNPs among various tissues (Figure 5b).

**Figure 5.**
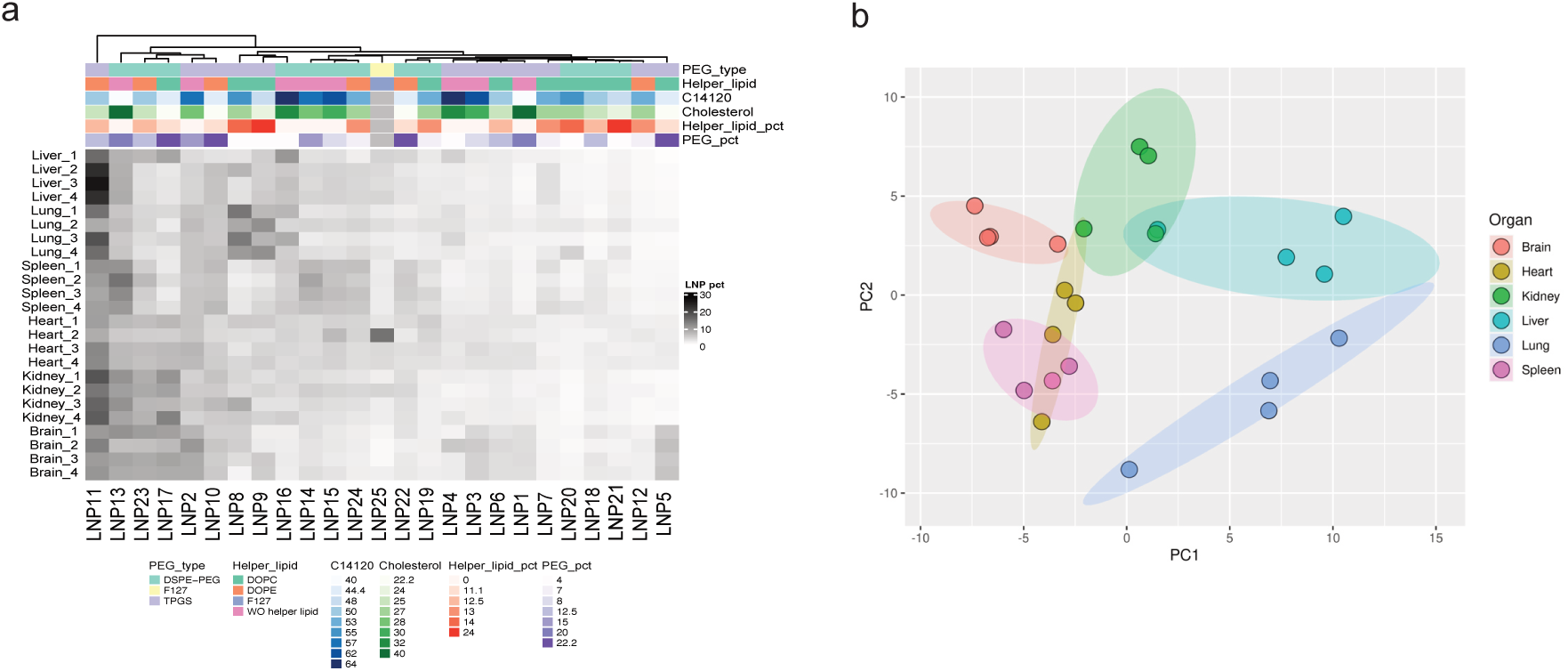
In vivo screening of LNP accumulation through sequencing. a) Heatmap showing the normalized DNA counts in various organs measured at 24 hours after i.v. injection of LNPs. (b) Principal component analysis (PCA) of normalized DNA counts showing the separation of different tissue clusters.

To assess the circulation characteristics of the individual LNPs, the relative distribution of DNA barcodes was analysed after 30 and 60 min in vivo circulation time. Heatmap analysis (Figure S8) suggested no significant differences in relative LNP levels. Although LNP11, LNP2 and LNP13 showed slightly higher read counts, the overall variation among formulations was minimal.

Moreover, H&E staining of liver and lung tissues from mice treated with the LNP library exhibited similar tissue structure to the negative control group treated with PBS. (Figure S9).

In summary, the barcoded and Luc-mRNA-containing LNPs exhibited strong and sustained bioluminescence mainly in liver and lung. Analysis of barcode sequences from various organs effectively revealed varying LNP accumulation patterns and allowed the identification of potential tissue-specific LNPs with targeting potential across different tissues (summarized in Table 2).

**Table 2.**
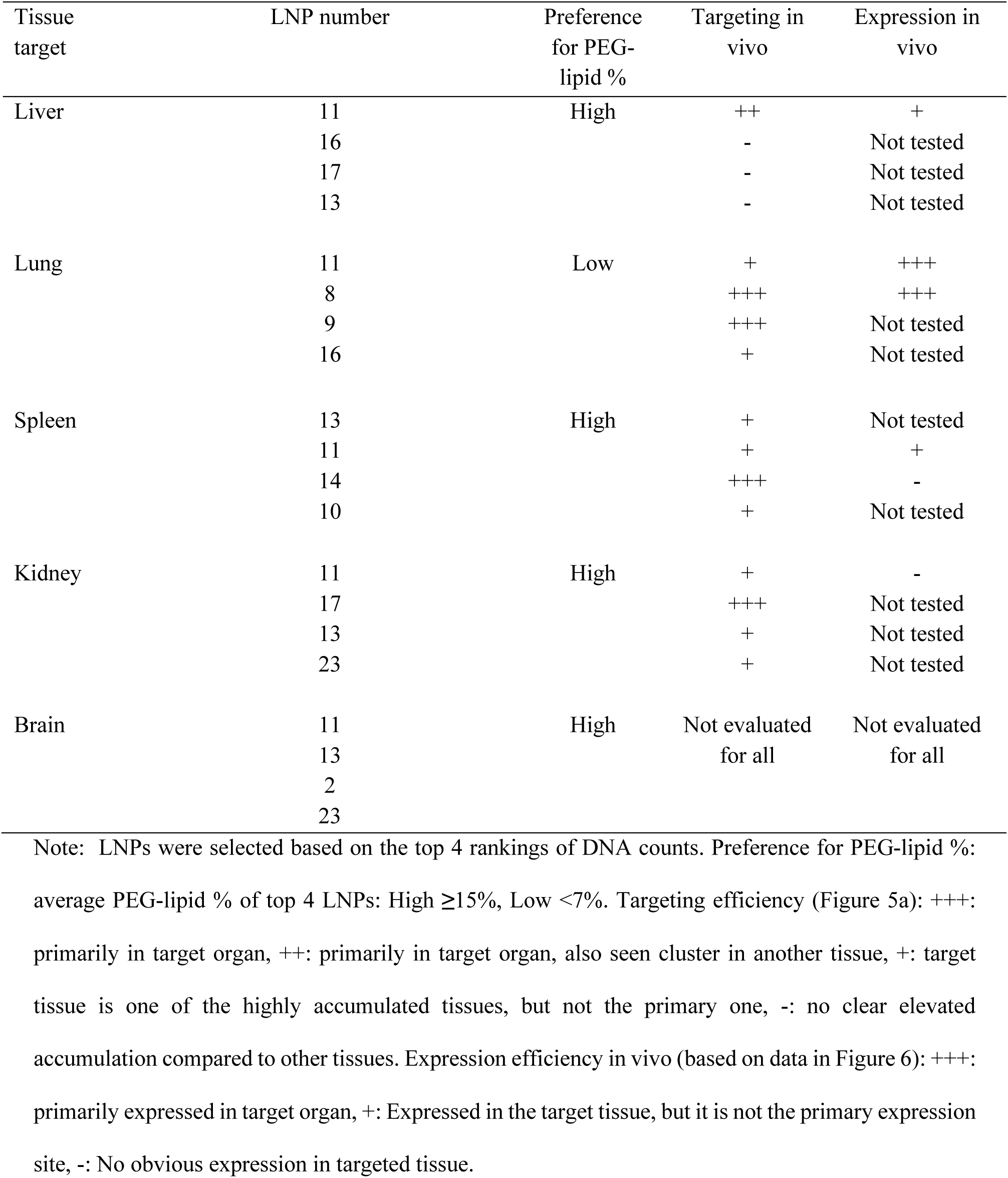
Summary of organ accumulation of LNPs, PEG-lipid% preferences, targeting specificity and in vivo mRNA expression specificity.

| Tissue target | LNP number | Preference for PEG-lipid % | Targeting in vivo | Expression in vivo |
| --- | --- | --- | --- | --- |
| Liver | 11 | High | ++ | + |
|  | 16 |  | - | Not tested |
|  | 17 |  | - | Not tested |
|  | 13 |  | - | Not tested |
| Lung | 11 | Low | + | +++ |
|  | 8 |  | +++ | +++ |
|  | 9 |  | +++ | Not tested |
|  | 16 |  | + | Not tested |
| Spleen | 13 | High | + | Not tested |
|  | 11 |  | + | + |
|  | 14 |  | +++ | - |
|  | 10 |  | + | Not tested |
| Kidney | 11 | High | + | - |
|  | 17 |  | +++ | Not tested |
|  | 13 |  | + | Not tested |
|  | 23 |  | + | Not tested |
| Brain | 11 | High | Not evaluated for all | Not evaluated for all |
|  | 13 |  |  |  |
|  | 2 |  |  |  |
|  | 23 |  |  |  |

Note: LNPs were selected based on the top 4 rankings of DNA counts. Preference for PEG-lipid %: average PEG-lipid % of top 4 LNPs: High $\geq 15\%$ , Low $< 7\%$ . Targeting efficiency (Figure 5a): +++: primarily in target organ, ++: primarily in target organ, also seen cluster in another tissue, +: target tissue is one of the highly accumulated tissues, but not the primary one, -: no clear elevated accumulation compared to other tissues. Expression efficiency in vivo (based on data in Figure 6): +++: primarily expressed in target organ, +: Expressed in the target tissue, but it is not the primary expression site, -: No obvious expression in targeted tissue.

### In vivo validation of LNP candidates

Based on the observed LNP signatures (Figure 5a, S6), we selected LNP8, LNP11 and LNP14 as the most promising candidates for lung, liver, and spleen targeting, respectively, and employed them individually to validate organ-specific functional delivery of Luc-mRNA at 6 hours after intravenous administration.

For LNP8, notable expression in lung tissue confirmed this targeting, while no significant expression compared to the control samples was observed in other organs (Figure 6a, d). LNP11 yielded, as expected, expression in the liver, but even higher expression in the lung and spleen (Figure 6b, d). Lastly, LNP14, considered as a candidate for spleen delivery, exhibited negligible expression within the spleen but strong expression within the liver and lung; however, statistical significance of the latter was not attained due to substantial variations. (Figure 6c, d). The specificity of the LNP candidates were further validated by analysing the luminescence ratios between the lung (or spleen) and liver. LNP8 and LNP11 exhibited luminescence intensities approximately 2-fold and 5-fold higher in the lung compared to the liver, respectively, while exhibited comparable expression levels between the spleen and liver. In contrast, LNP14 demonstrated similar luminescence intensities in the liver and lung, with a ratio close to 1, whereas spleen expression was markedly lower (Figure 6e).

**Figure 6.**
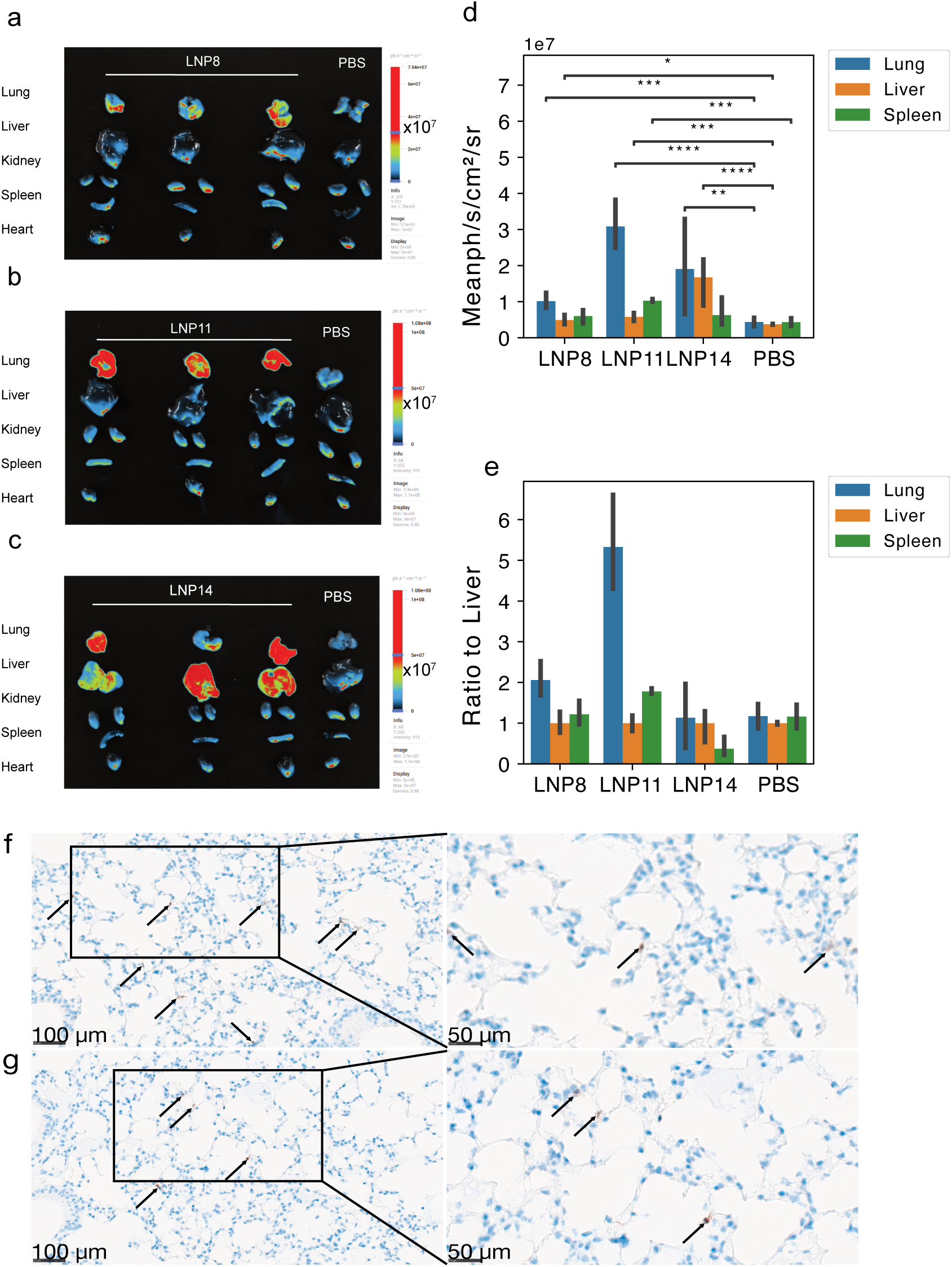
Luciferase expression in various organs using selected organ-targeted LNP candidates, using lung-targeting LNP8 (a), liver targeting LNP11 (b) and spleen targeting LNP14 (c). IVIS image is shown in a-c and average intensity of region of interest (ROI) are shown in d. The luminescence ratio of Lung or Spleen to each individual liver signal is shown in e. Data are presented as Mean+-SD (n =3). * *p < 0.05*, ** *p < 0.01*, *** *p < 0.001*. Significance was determined by independent *t*-test. (f) and (g) Representative images of mCherry immunohistochemical staining in normal lung tissue sections from two individually treated mice. Positive mCherry signals, visualized by brown 3,3′-Diaminobenzidine (DAB) chromogen and indicated by arrows, are localized to scattered pneumocytes along the alveolar surface. Sections were counterstained with haematoxylin to visualize nuclei. Images are shown at both overview and high magnification. Scale bars indicated in the lower left corner are 100 µm (overview) and 50 µm (high magnification).

To further validate lung tropism of LNP8 demonstrated by the increased luciferase expression specifically in lung tissue, and identify the targeted cell populations, we encapsulated mCherry mRNA and administered i.v. to mice. Immunohistochemical analysis of lung tissues revealed mCherry-positive staining in scattered alveolar epithelial cells along the alveolar surface (Figure 6f, g), which was absent in PBS-treated controls (Figure S10). These findings indicate that LNP8 reached the pulmonary parenchyma from the circulation and enabled functional mRNA expression within pneumocytes.

## Discussion

Identification of LNPs with unique target specificity and efficiency is critical for further development of RNA-based therapeutic applications. Additionally, elucidating the functional implications of varying the type and ratio of lipids in terms of their biodistribution and expression profiles is crucial for the rational design of LNPs. Compared to conventional approaches where LNPs are designed and evaluated individually, high-throughput multiplexed barcode screening methodologies offer advantages by substantial reduction in animal consumption and experimental workload for screening large numbers of constituents. Here we employed both approaches for screening of 25 C14120-based LNPs, enabling us to identify highly potent LNPs for mRNA expression in vitro (Table 1) and tissue targeting capacity in vivo (Table 2).

Characterization of the LNPs demonstrated well-defined physicochemical properties in terms of size, polydispersity, zeta potential and mRNA encapsulation efficiency. These properties were influenced to varying extents by the chemical composition of the LNPs, with a determining influence of the PEG lipid, which affected size, polydispersity and overall functional delivery of mRNA. For example, in in vitro experiments, the PEG-lipid percentage showed opposing effects on mRNA expression in human and mouse liver cell lines or human cell lines from different tissues. In the codelivery experiment, LNPs exhibited significantly higher cellular uptake levels in AML12 compared to HEK293-H. This is likely due to the distinct cell origin with different metabolic cellular characteristics, in accordance with previous reports showing that cellular uptake can vary dramatically with up to 50 times between cell types ^39–41^. Additionally, factors such as difference in mRNA degradation rate between cell lines may contribute to this disparity. Importantly, the ionizable lipid within LNPs become positively charged in the acidic endosomal environment, which help protect the encapsulated DNA barcode. The degradation is more likely to occur either after endosomal escape or in lysosome. The mRNA and DNA barcode that achieved endosomal escape is generally considered to be very low (1-2%), and may therefore have minimal impact on the overall uptake levels. Furthermore, the codelivery of mRNA and DNA barcode experiment revealed that while cellular uptake is critical for RNA delivery, the bias between cellular uptake and mRNA expression levels indicates that uptake alone does not determine the expression efficiency, suggesting that factors such as endosomal escape, which may vary based on LNP formulation and cell origin, plays a significant role. This cell line-specific behaviour and discrepancy underlines the importance of comprehensive evaluation of LNPs, which was further highlighted by comparison to in vivo delivery efficiencies. Here, we observed significant discrepancies when selecting the best-performing candidates for in vitro and in vivo delivery. Specifically, among the top performing LNPs for uptake in mouse liver AML12 cells (LNP4, LNP10-12 and LNP23, Figure S4c-d), only LNP11 demonstrated high accumulation in mouse liver, while the rest exhibited minimal accumulation in mouse liver tissue, concerning the common practice to use best-performing LNPs for mRNA expression in vitro for in vivo applications. For example, the top LNPs in human liver Hep G2 cells (LNP2, LNP5, Figure S3b) and mouse liver AML12 cells (LNP4, Figure S4c, S3c) exhibited minimal accumulation in mouse liver in vivo (Figure 5a). Similarly, LNP16, LNP19, LNP24 (with low PEG %), demonstrated pronounced potency in expression in bEnd.3 (Figure S3f), but failed to display as efficient accumulation in mouse brain (Figure S6f). Conversely, LNP5 (high PEG %) exhibited greater accumulation within brain tissues. This confirms that in vitro studies are not reliable predictors of in vivo performance and biodistribution^42,43^.

In vivo administration of the combined barcoded LNP library containing Luc-mRNA resulted in robust expression in the liver at 6 hours, with a shift towards more pronounced expression in the lung after 24 hours. This result was expected based on previous studies showing that mRNA delivered via LNPs accumulate primarily in the liver, and to a varied extent and timing in organs such as spleen, lungs and kidneys based on LNP properties ^44,45^.

Our investigation of the biodistribution in relation to lipid percentages present in the formulations, highlights PEG-lipid and helper lipid as determining factors (Figure S7). However, discerning a direct correlation between tissue targeting and PEG-lipid percentages (as well as other lipid components) is challenging. This difficulty arises from the predominant representation of the top-performing LNP formulations within the dataset, particularly LNP11, which shows the most prominent accumulation in liver and elevated levels across various other organ types.

The positive effect of DOPE incorporation on liver uptake is clear and supported by a previous report which suggested that DOPE interacts with apolipoprotein E (ApoE), exerting affinity for low-density lipoprotein (LDL) receptor highly expressed by hepatocytes^46^. LNP11 is not the exclusive formulation within the library containing DOPE; Hence, the content of other components also influences liver specificity.

Regarding the coating lipid, Dilliard et al. has proposed a mechanism that involves PEG-lipid desorption from the LNP surface followed by their association with specific serum proteins that dictate tissue targeting. This association of these organ-specific serum proteins strongly correlates with the charged lipid and ratios within the formulation ^47,48^. LNP11 contains TPGS, which exhibits shorter PEG chain and smaller lipid tail (vitamin E) compared to DSPE-PEG, leading a higher propensity for TPGS desorption, potentially enhancing the interaction with ApoE thereby augmenting the liver targeting.

A closer examination of the 4 most abundant LNPs accumulating in various organs reveals a preference for high percentage of PEG-lipid within formulations in spleen, kidney, heart and brain tissues, while in the context of lung delivery, 3 out of the top 4 LNPs are formulated with low percentage (4%) of PEG-lipid (Figure S6).

Lastly, we aimed to complement the biodistribution analysis with mRNA expression analysis for individual promising LNP candidates with preference for lung (LNP8), liver (LNP11) and spleen (LNP14) targeting. The detection of mCherry protein in alveolar epithelial cells after systemic administration indicates that LNP8 effectively reached the lung parenchyma via the bloodstream, crossed the pulmonary microvascular barrier and enabled functional mRNA expression within pneumocytes, supporting its potential for future respiratory applications. However, we did not find a clear correlation between accumulation and expression for all candidates (Figure 6a-e). This discrepancy can be due to several factors. Apart from homing into specific tissues, LNP also must mediate cellular uptake, endosomal escape and release of the mRNA before engaging the ribosomes, in which the efficiency of uptake and intracellular trafficking of different LNPs varies significantly^49,50^. Additionally, the stability of mRNA loaded in different LNPs and their translation machinery availability in different cells impact expression efficiency^51^. Variations in the immune response triggered by the LNPs can further lead to the inhibition of translation and degradation of the expressed protein^52^. Furthermore, microenvironmental factors, including enzymatic activity, pH and cellular conditions, can potentially affect both the stability and mRNA expression efficiency^53^. These factors collectively determine the complex relationship between biodistribution and mRNA expression. One limitation of our approach is that the barcode tracking solely reveals accumulation levels rather than functional mRNA expression. However, the observed capacity of certain LNPs to target specific organs can be used as first guideline to select optimal lipid composition for delivery.

In summary, we have identified LNP compositions optimized for in vitro mRNA expression in a panel of cell lines and for targeting of specific tissues in vivo. The results provide guidelines (summarized in Table 1 and 2) for designing LNPs with improved functional delivery of mRNA in vitro and in vivo.

## Materials and Methods

### Materials

Pluronic F127, d-α-tocopheryl polyethyleneglycol 1000 succinate (TPGS), 1,2-epoxytetradecane (ETD), 2,2’-ethylenedioxybisethylamine (EOBA) and cholesterol were purchased from Sigma-Aldrich. 2-distearoyl-sn-glycero-3-phosphoethanolamine-N-[methoxy(polyethylene glycol)-2000] (DSPE– PEG-2000), 1,2-dioleoyl-sn-glycero-3-phosphoethanolamine (DOPE) and 1,2-dioleoyl-sn-glycero-3-phosphocholine (DOPC) were purchased from Avanti Polar Lipids. CleanCap mCherry mRNA and luciferase (Luc) mRNA were purchased from TriLink Biotechnologies.

### Synthesis of C14120

The lipid-like material C14120 was synthesized by a ring-opening reaction between ETD and EOBA at a molar ratio of 3:1. The vial was sealed, and the reaction was conducted at 90°C with stirring for 2.5 days in the dark. The crude product was purified by chromatography on silica gel with gradient elution from dichloromethane (CH2Cl2) to the CH2Cl2/MeOH/NH4OH (75:22:3), yielding a viscous pale-yellow oil. The purified product was verified with MALDI-TOF MS recorded on a Bruker BIFLEX III spectrometer and 1H NMR spectroscopy and stored at -20°C until further use.

### Preparation of nanoparticles

A library of 25 lipid nanoparticles (LNPs) was synthesized by nanoprecipitation. Briefly, C14120, cholesterol, DOPC, DOPE, TPGS, F127, and DSPE-PEG were dissolved in 100% ethanol and combined at different ratios (Figure 2a). Acetate buffer (200 mM, pH 5.5) was quickly injected into the lipid mixture while stirring to self-assemble into empty LNPs. Subsequent dialysis (3.5 KDa cutoff, Biodesign™) of the LNP against acetate buffer (200 mM, pH 5.5) was performed to remove residual ethanol. For encapsulation, mRNA was mixed with LNPs individually at an N/P ratio of 12 and incubated at room temperature for 10 minutes (min).

### DNA barcoding of LNPs

58 nucleotide (nt)-long ssDNA sequences were purchased from Integrated DNA Technologies (IDT). Each individual LNP was prepared to carry its own DNA barcode (Fig.1). To encapsulate the DNA barcode, mRNA and DNA barcodes were mixed at weight ratio of 10:1. The resulting mixture was further mixed with empty nanoparticles to generate LNPs containing mRNA and its unique barcode. The DNA barcode library and primer sequences are described in Table S1-S2.

### Characterization of LNPs

The size and zeta potential of the LNPs were measured by Dynamic Light Scattering (DLS) at 25°C on a Zetasizer Nano ZS (Malvern Instruments, Malvern, UK). The quantification of mRNA encapsulation efficiency was determined by RiboGreen Reagent (Invitrogen) according to the manufacturer’s instructions. Briefly, the LNP/mRNA solution was mixed with RiboGreen solution (1:100 in TE buffer) at a volume ratio of 1:1 and incubated for 5 min at room temperature. The fluorescent intensity was measured by FLUOstar OPTIMA (Moritex BioScience) at excitation and emission wavelengths of 480 nm and 520 nm, respectively.

### Cell culture

Human retinal pigment epithelial cell line ARPE-19 (ATCC Manassas, VA, USA), mouse brain endothelial cell line bEnd.3 (ATCC Manassas, VA, USA), human hepatoma cell line Hep G2 (ATCC Manassas, VA, USA), mouse hepatocytes AML12 (ATCC Manassas, VA, USA) and human embryonic kidney cell line HEK293-H (Thermo Scientific Waltham, MA, USA) were maintained in cell culture medium supplemented with 10% fetal bovine serum and 1% penicillin-streptomycin at 37°C, 5% CO_2_ and 100% humidity. ARPE-19 and AML12 cells were maintained in Dulbecco’s Modified Eagle’s Medium F12 (DMEM F12, ThermoFisher, Waltham, MA, USA), while Hep G2, bEnd.3 and HEK293-H cells were maintained in Dulbecco’s Modified Eagle’s Medium (DMEM, Thermo Scientific, Waltham, MA, USA).

### In vitro mRNA expression studies

ARPE-19 (3×10^4^/well), bEnd.3 (5×10^4^/well), Hep G2 (2×10^5^/well), AML12 (2.5×10^4^/well) and HEK293-H (2×10^5^/well) cells were seeded in 48-well plates. After overnight incubation, the medium was replaced with 300 μL fresh medium, and the cells were transfected with LNPs encapsulating mCherry mRNA at a final dose of 100 ng/well. For comparison, 100 ng of RNA/well were transfected using Lipofectamine 2000 (Thermo Scientific) according to the manufacturer’s instructions. After 24 hours (h), the cells were imaged by fluorescent microscope (OLYMPUX IX71) and subsequently detached using 0.05% Trypsin-EDTA and analyzed by NovoCyte flow cytometer (ACEA Biosciences).

### Codelivery of mRNA and barcode in vitro

HEK293-H (2×10^5^/well) and AML12 cells (3×10^4^/well) were seeded using the same method as above co-transfected both mRNA encoding EYFP and barcode (Barcode-ATTO647) encapsulated in individual LNPs from LNP1-24 and compared with the commercialized formulation, SM102 (Moderna). A final dose of mRNA and Barcode of 100ng and 10ng/well was used for the codelivery. After 24 hours incubation, the cells were detached using 0.05% Trypsin-EDTA and analysed by NovoCyte flow cytometer (ACEA Biosciences).

### Particle mixing test

Two LNP candidates (LNP19 and LNP25) were selected to assess the possibility of particle mixing. Briefly, the LNPs were individually mixed with 400 ng of Cy3-labeled siRNA (Cy3-siRNA), Cy5-labeled siRNA (Cy5-siRNA), or co-encapsulated with both Cy3-siRNA and Cy5-siRNA at an N/P ratio of 12. After incubation for 10 min at room temperature, the Cy3-siRNA containing LNPs were mixed thoroughly with Cy5-siRNA containing LNPs. Particle content mixing was evaluated by Fluorescence Resonance Energy Transfer (FRET) between donor Cy3 to acceptor Cy5 (Cy5/Cy3 ratio) under laser excitation for Cy3 at 532 nm, emission for Cy3 (570 nm) and Cy5(670 nm) was recorded using an Amersham^TM^ Typhoon scanner. The sense (5′-GACGUAAACGGCCACAAGUTC-NH2-3′) and antisense (5′-ACUUGUGGCCGUUUACGUCGC-3′) strands of the siRNA were purchased from Ribotask (Denmark), and the sense strand was labelled at the 3’ end with Cy3-NHS or Cy5-NHS through click chemistry. The siRNAs were annealed in 10 mM Tris, pH 8.0, 20 mM NaCl (Figure S5b).

### In vivo delivery of LNP library containing mRNA and DNA barcodes

C57BL/6J 6-8 week old mice were purchased from Janvier Labs. To screen the nanoparticles in vivo, 8 mice were administered the barcoded LNP library through tail vein injection (i.v.) at mRNA (Luc-mRNA) dose of 1 mg/kg with the weight ratio between mRNA and Barcode of 10:1^25^. Subsequently, the mice were intraperitoneal injected (i.p.) with D-luciferin substrate at 150 mg/kg and scanned after 10 min with an IVIS 200 imaging system (Xenogen, Caliper Life Sciences, Hopkinton, MA, USA) after 6 h and 24 h. The mice were sacrificed, and organs were harvested, scanned with IVIS, and collected for further studies.

### Isolation and amplification of barcode sequences for NGS sequencing

Tissues were minced and lysed with RIPA lysis buffer and subsequently incubated at 55°C while shaking overnight. The crude samples were then centrifuged at 14000 g x 10 min at room temperature. The supernatant was collected and further purified with ZYMO ssDNA/RNA Clean & Concentrate columns (D7011)^33^. The plasma samples were lysed with RIPA lysis buffer and purified directly with ZYMO ssDNA/RNA Clean & Concentrate columns. Each barcode pool was amplified by PCR using Phusion High-Fidelity DNA polymerase kit (Thermo Scientific) and 4 μL 5x HF Phusion buffer, 0.4 μL 10 mM dNTPs (Thermo Scientific), 2 μL template oligonucleotide pool, 0.2 μL Phusion polymerase, 2 μL DMSO, 0.25 μL universal forward primer (10 μM), 0.25 μL Reverse primer (10 μM), 10.9 μL H_2_O. PCR conditions included initial denaturation at 98°C for 1.5min and 15-25 cycles of 98°C for 15 seconds (s), 60°C for 15 s, and 68°C for 30 s. The barcodes from liver, lung, spleen, kidney and plasma were PCR amplified for 15 cycles, while barcodes from heart and brain were amplified for 25 cycles.

PCR products were further purified by AMPure beads (Beckman Coulter) by mixing 1:1 (volume ratio), the supernatant was collected and mixed 2:1 (volume ratio) with AMPure beads. The beads were collected and washed twice with 80% ethanol before the product was eluted in 15 μL RNase free water. Prior to sequencing, the purity and concentration of the samples were assessed using the Agilent bioanalyser 2100 (Figure S11). The samples were pooled equimolar (15 fmol) and submitted to the MOMA NGS core center at Aarhus University Hospital and sequenced on an Illumina MiSeq platform with paired-end 75 sequencing, resulting in 25 million reads. The data generated from Illumina sequencing was processed by Omiics (Aarhus, Denmark), where the barcodes were recognized using the pattern matching by partial ratio function implemented in the RipidFuzz python package. In the sequences, a partial ratio of 87 (allowing for 1 mismatch) was used for both the 8 nt barcode and index primer for pattern matching, which generated an output of counts for all sequences in each sample. The raw counts were normalized to percentage through dividing each raw count by the total counts within each sample.

Furthermore, the liver and lung tissues were fixed in 4% PFA, embedded in paraffin, and cut into 4 μm-thick sections. Histologic investigations of H&E staining were carried out using Haematoxylin Solution (Mayer’s, Modified) and Eosin Y solution, Alcoholic. The stained sections were imaged by an Olympus IX71 microscope (Olympus, US) at 20x and 10x magnification with bright field (BF).

### Validation experiment in mice based on LNP candidates

LNP candidates encapsulating Luc-mRNA were administered i.v. to 6-8 weeks old C57BL/6J mice at a dose of 10 μg/mouse (0.5 mg/kg). Subsequently, D-luciferin substrate was administered i.p. at 150 mg/kg at 6h. After 10 min the mice were sacrificed, and all organs were isolated and scanned with a Newton 7.0 optical imaging system (Vilber, France).

LNP8 was subsequently selected for mCherry mRNA delivery to visualize expression in lung tissues. LNP8 encapsulating mCherry mRNA was administered i.v. to 6-8-week old C57BL/6J mice at a dose of 0.75 mg/kg (n = 3); a PBS-treated group served as the negative control (n = 3). 6 h post-injection, mice were sacrificed, and lung tissues were isolated and fixed in 10% neutral buffered formalin for immunohistochemistry (IHC).

IHC was performed on a Ventana Discovery Ultra instrument (Ventana Medical System). The primary antibody used was rabbit monoclonal antibody detecting mCherry: (EPR20579, Abcam; RRID: AB_213511) diluted 1:100. After deparaffination and rehydration of the sections, antigen retrieval was performed using CC2 buffer (Ventana Medical System) at 99°C for 30 min. Slides were treated with inhibitors of endogenous peroxidase followed by incubation with primary antibody for 32 min at 36°C. Signal amplification was performed using DISCOVERY OmniMap antiRB HRP (Ventana Medical System). Visualization was done using OptiView DAB IHC Detection Kit 760/700 (Ventana Medical System) and hematoxylin was used as counterstain. Data were collected using a Hamamatsu NanoZoomer XR digital slide scanner and analysed using NDPview 2 software version 2.9.29 for Mac (OSX).

### Data analysis

The linear regression was performed using lmplot function in seaborn package in Python (Python 3.9.7). The heatmap was plotted by Omiics (Aarhus, Denmark) using R. PCA analysis was performed using prcomp function and visualized using ggplot2 package in R^54^. The data points for FACS data were measured and exported by FCMeasurement function from FlowCytometryTools package (http://eyurtsev.github.io/FlowCytometryTools) in Python, and the histogram overlay for flow cytometry data was conducted by joyplot function in joypy package in Python^55^. All the barplots, scatter plots and lineplots in the study were conducted using Python seaborn package. The significance was determined by independent t-test. Scripts are available upon reasonable request.

## Acknowledgements

The work is supported by Novo Nordisk Foundation grant NNF23OC0081177 (RNA-META). C.V, J.S and S.F were funded by the Danish National Research Foundation (Centre for Cellular Signal Patterns, DNRF135).

## Author Contributions

P.S and J.K conceived the project and planned the research. P.S carried out the experiments other than the sequencing analysis. J.S helped in sample preparation for sequencing and S.F helped in data analysis for the PCA plot. C.V helped in the validation animal experiment. M.G helped extensively in helpful writing and revision. P.S analyzed the data and wrote the original draft with the extensive help in revision from J.K.

## Declaration of interests

The authors declare no competing interests.

## Data Availability

Research materials and data are available upon reasonable request to the corresponding author.

## Notes

### Competing Interest Statement

The authors have declared no competing interest.

