## supporting information for "Optimizing C14120-based LNPs for in vitro and in vivo mRNA delivery"

**Supplemental Information**

Table S1. The design and sequences of DNA barcodes in the library

G*G*A*TGTGCTGCGAGAAGGCTAGANNNNTGATATTGTCTAGCCTTCTCGTGTGCA GA*C*T

Red = phosphorothioate linkages (*) act to increase resistance to exonucleases. Green = Universal primer binding sites allow for amplification from cells/tissues and linkage to next generation sequence adapters Deep red = 8nt Barcode. This 8nt sequence is referenced to identify nanoparticle composition and track nanoparticle distribution. Light Blue= 4nt Random nucleotide region used to minimize PCR bias.

| LNP | Barcode sequence 5'-3' |
|---|---|
| LNP1 | GGATGTGCTGCGAGAAGGCTAGANNNNTGATATTGTCTAGCCTTCTCGTGTGCAGACT |
| LNP2 | GGATGTGCTGCGAGAAGGCTAGANNNNGCGAGTATTCTAGCCTTCTCGTGTGCAGACT |
| LNP3 | GGATGTGCTGCGAGAAGGCTAGANNNNGATCTACCTCTAGCCTTCTCGTGTGCAGACT |
| LNP4 | GGATGTGCTGCGAGAAGGCTAGANNNNATGAGATGTCTAGCCTTCTCGTGTGCAGACT |
| LNP5 | GGATGTGCTGCGAGAAGGCTAGANNNNAGCATGCGTCTAGCCTTCTCGTGTGCAGACT |
| LNP6 | GGATGTGCTGCGAGAAGGCTAGANNNNTACCTGCTTCTAGCCTTCTCGTGTGCAGACT |
| LNP7 | GGATGTGCTGCGAGAAGGCTAGANNNNCTCCTTCGTCTAGCCTTCTCGTGTGCAGACT |
| LNP8 | GGATGTGCTGCGAGAAGGCTAGANNNNGCAGGACTTCTAGCCTTCTCGTGTGCAGACT |
| LNP9 | GGATGTGCTGCGAGAAGGCTAGANNNNCGCCTATTTCTAGCCTTCTCGTGTGCAGACT |
| LNP10 | GGATGTGCTGCGAGAAGGCTAGANNNNTCCTAAGATCTAGCCTTCTCGTGTGCAGACT |
| LNP11 | GGATGTGCTGCGAGAAGGCTAGANNNNCAAGAAGGTCTAGCCTTCTCGTGTGCAGACT |
| LNP12 | GGATGTGCTGCGAGAAGGCTAGANNNNTAGATCCGTCTAGCCTTCTCGTGTGCAGACT |
| LNP13 | GGATGTGCTGCGAGAAGGCTAGANNNNTAAGATGATCTAGCCTTCTCGTGTGCAGACT |
| LNP14 | GGATGTGCTGCGAGAAGGCTAGANNNNTAACCGAATCTAGCCTTCTCGTGTGCAGACT |
| LNP15 | GGATGTGCTGCGAGAAGGCTAGANNNNACGTCGAATCTAGCCTTCTCGTGTGCAGACT |
| LNP16 | GGATGTGCTGCGAGAAGGCTAGANNNNTTGCAACTTCTAGCCTTCTCGTGTGCAGACT |
| LNP17 | GGATGTGCTGCGAGAAGGCTAGANNNNTATGCCTTTCTAGCCTTCTCGTGTGCAGACT |
| LNP18 | GGATGTGCTGCGAGAAGGCTAGANNNNGTCTCCGTTCTAGCCTTCTCGTGTGCAGACT |
| LNP19 | GGATGTGCTGCGAGAAGGCTAGANNNNAGTCCGGTTCTAGCCTTCTCGTGTGCAGACT |
| LNP20 | GGATGTGCTGCGAGAAGGCTAGANNNNATCGTCTATCTAGCCTTCTCGTGTGCAGACT |

| | |
|---|---|
| LNP21 | GGATGTGCTGCGAGAAGGCTAGANNNNCAATCCGTTCTAGCCTTCTCGTGTGCAGACT |
| LNP22 | GGATGTGCTGCGAGAAGGCTAGANNNNGTCCGTTATCTAGCCTTCTCGTGTGCAGACT |
| LNP23 | GGATGTGCTGCGAGAAGGCTAGANNNNCATAATAGTCTAGCCTTCTCGTGTGCAGACT |
| LNP24 | GGATGTGCTGCGAGAAGGCTAGANNNNATTCGAGATCTAGCCTTCTCGTGTGCAGACT |
| LNP25 | GGATGTGCTGCGAGAAGGCTAGANNNNGTTAGTCATCTAGCCTTCTCGTGTGCAGACT |

Table S2. The adapter primer sequences for PCR amplification.

| Mouse tissue label | Universal primer | Sequence | Index primer | Sequence |
|---|---|---|---|---|
| Liver 1 | U1 | 5′-AATGATACGGCGACCACCGAGATCTACACTATAGCCTGGATGTGCTGCGAGAAGGCTAGA-3’ | In1 | 5′-CAAGCAGAAGACGGCATACGAGATATTACTGCAGTCTGCACACGAGAAGGCTAGA-3’ |
| Liver 2 | U1 | 5′-AATGATACGGCGACCACCGAGATCTACACTATAGCCTGGATGTGCTGCGAGAAGGCTAGA-3’ | In2 | 5′-CAAGCAGAAGACGGCATACGAGATTCCGGAGAAGTCTGCACACGAGAAGGCTAGA-3’ |
| Liver 3 | U1 | 5′-AATGATACGGCGACCACCGAGATCTACACTATAGCCTGGATGTGCTGCGAGAAGGCTAGA-3’ | In3 | 5′-CAAGCAGAAGACGGCATACGAGATCGCTCATTAGTCTGCACACGAGAAGGCTAGA-3’ |
| Liver 4 | U1 | 5′-AATGATACGGCGACCACCGAGATCTACACTATAGCCTGGATGTGCTGCGAGAAGGCTAGA-3’ | In4 | 5′-CAAGCAGAAGACGGCATACGAGATGAGATTCCAGTCTGCACACGAGAAGGCTAGA-3’ |
| Lung 1 | U1 | 5′-AATGATACGGCGACCACCGAGATCTACACTATAGCCTGGATGTGCTGCGAGAAGGCTAGA-3’ | In5 | 5′-CAAGCAGAAGACGGCATACGAGATATTCAGAAAGTCTGCACACGAGAAGGCTAGA-3’ |
| Lung 2 | U1 | 5′-AATGATACGGCGACCACCGAGATCTACACTATAGCCTGGATGTGCTGCGAGAAGGCTAGA-3’ | In6 | 5′-CAAGCAGAAGACGGCATACGAGATGAATTCGTAGTCTGCACACGAGAAGGCTAGA-3’ |
| Lung 3 | U1 | 5′-AATGATACGGCGACCACCGAGATCTACACTATAGCCTGGATGTGCTGCGAGAAGGCTAGA-3’ | In7 | 5′-CAAGCAGAAGACGGCATACGAGATCTGAAGCTAGTCTGCACACGAGAAGGCTAGA-3’ |
| Lung 4 | U1 | 5′-AATGATACGGCGACCACCGAGATCTACACTATAGCCTGGATGTGCTGCGAGAAGGCTAGA-3’ | In8 | 5′-CAAGCAGAAGACGGCATACGAGATTAATGCGCAGTCTGCACACGAGAAGGCTAGA-3’ |
| Spleen 1 | U1 | 5′-AATGATACGGCGACCACCGAGATCTACACTATAGCCTGGATGTGCTGCGAGAAGGCTAGA-3’ | In9 | 5′-CAAGCAGAAGACGGCATACGAGATCGGCTATGAGTCTGCACACGAGAAGGCTAGA-3’ |
| Spleen 2 | U1 | 5′-AATGATACGGCGACCACCGAGATCTACACTATAGCCTGGATGTGCTGCGAGAAGGCTAGA-3’ | In10 | 5′-CAAGCAGAAGACGGCATACGAGATTCCGCGAAAGTCTGCACACGAGAAGGCTAGA-3’ |
| Spleen 3 | U1 | 5′-AATGATACGGCGACCACCGAGATCTACACTATAGCCTGGATGTGCTGCGAGAAGGCTAGA-3’ | In11 | 5′-CAAGCAGAAGACGGCATACGAGATTCTCGCGCAGTCTGCACACGAGAAGGCTAGA-3’ |
| Spleen 4 | U1 | 5′-AATGATACGGCGACCACCGAGATCTACACTATAGCCTGGATGTGCTGCGAGAAGGCTAGA-3’ | In12 | 5′-CAAGCAGAAGACGGCATACGAGATAGCGATAGAGTCTGCACACGAGAAGGCTAGA-3’ |
| Heart 1 | U2 | 5′-AATGATACGGCGACCACCGAGATCTACACATAGAGGCGGATGTGCTGCGAGAAGGCTAGA-3’ | In13 | 5′-CAAGCAGAAGACGGCATACGAGATCCTATCCTAGTCTGCACACGAGAAGGCTAGA-3’ |
| Heart 2 | U2 | 5′-AATGATACGGCGACCACCGAGATCTACACATAGAGGCGGATGTGCTGCGAGAAGGCTAGA-3’ | In14 | 5′-CAAGCAGAAGACGGCATACGAGATGGCTCTGAAGTCTGCACACGAGAAGGCTAGA-3’ |
| Heart 3 | U2 | 5′-AATGATACGGCGACCACCGAGATCTACACATAGAGGCGGATGTGCTGCGAGAAGGCTAGA-3’ | In15 | 5′-CAAGCAGAAGACGGCATACGAGATAGGCGAAGAGTCTGCACACGAGAAGGCTAGA-3’ |
| Heart 4 | U2 | 5′-AATGATACGGCGACCACCGAGATCTACACATAGAGGCGGATGTGCTGCGAGAAGGCTAGA-3’ | In16 | 5′-CAAGCAGAAGACGGCATACGAGATTAATCTTAAGTCTGCACACGAGAAGGCTAGA-3’ |
| Kidney 1 | U2 | 5′-AATGATACGGCGACCACCGAGATCTACACATAGAGGCGGATGTGCTGCGAGAAGGCTAGA-3’ | In17 | 5′-CAAGCAGAAGACGGCATACGAGATCAGGACGTAGTCTGCACACGAGAAGGCTAGA-3’ |
| Kidney 2 | U2 | 5′-AATGATACGGCGACCACCGAGATCTACACATAGAGGCGGATGTGCTGCGAGAAGGCTAGA-3’ | In18 | 5′-CAAGCAGAAGACGGCATACGAGATGTACTGACAGTCTGCACACGAGAAGGCTAGA-3’ |
| Kidney 3 | U2 | 5′-AATGATACGGCGACCACCGAGATCTACACATAGAGGCGGATGTGCTGCGAGAAGGCTAGA-3’ | In19 | 5′-CAAGCAGAAGACGGCATACGAGATACCCAGCAAGTCTGCACACGAGAAGGCTAGA-3’ |
| Kidney 4 | U2 | 5′-AATGATACGGCGACCACCGAGATCTACACATAGAGGCGGATGTGCTGCGAGAAGGCTAGA-3’ | In20 | 5′-CAAGCAGAAGACGGCATACGAGATAACCCCTCAGTCTGCACACGAGAAGGCTAGA-3’ |
| Brain 1 | U1 | 5′-AATGATACGGCGACCACCGAGATCTACACTATAGCCTGGATGTGCTGCGAGAAGGCTAGA-3’ | In21 | 5′-CAAGCAGAAGACGGCATACGAGATATCACGACAGTCTGCACACGAGAAGGCTAGA-3’ |
| Brain 2 | U1 | 5′-AATGATACGGCGACCACCGAGATCTACACTATAGCCTGGATGTGCTGCGAGAAGGCTAGA-3’ | In22 | 5′-CAAGCAGAAGACGGCATACGAGATACAGTGGTAGTCTGCACACGAGAAGGCTAGA-3’ |
| Brain 3 | U1 | 5′-AATGATACGGCGACCACCGAGATCTACACTATAGCCTGGATGTGCTGCGAGAAGGCTAGA-3’ | In23 | 5′-CAAGCAGAAGACGGCATACGAGATCAGATCCAAGTCTGCACACGAGAAGGCTAGA-3’ |
| Brain 4 | U1 | 5′-AATGATACGGCGACCACCGAGATCTACACTATAGCCTGGATGTGCTGCGAGAAGGCTAGA-3’ | In24 | 5′-CAAGCAGAAGACGGCATACGAGATACAAACGGAGTCTGCACACGAGAAGGCTAGA-3’ |

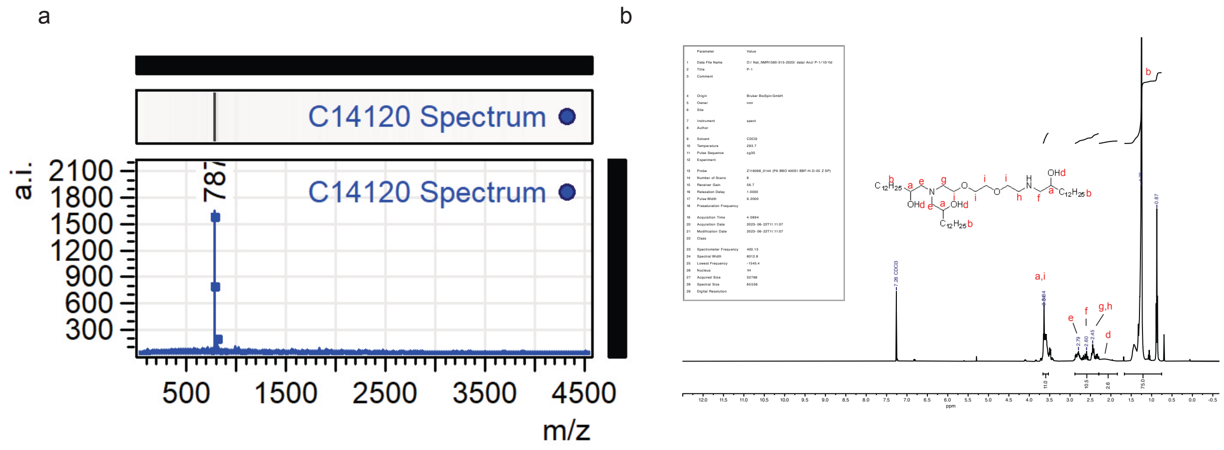


Figure S1. Structure of C14120 validated by (a) MALDI-TOF spectrum calculated with $[M+H]^+$ 786.3, found 787, and (b) $^1H$ NMR analysis with 3.68-3.50 (m, 11H), 3.00-2.26 (m, 10.5H), 2.23-1.90 (m, 2.6H), 1.71-0.77 (m, 75H).

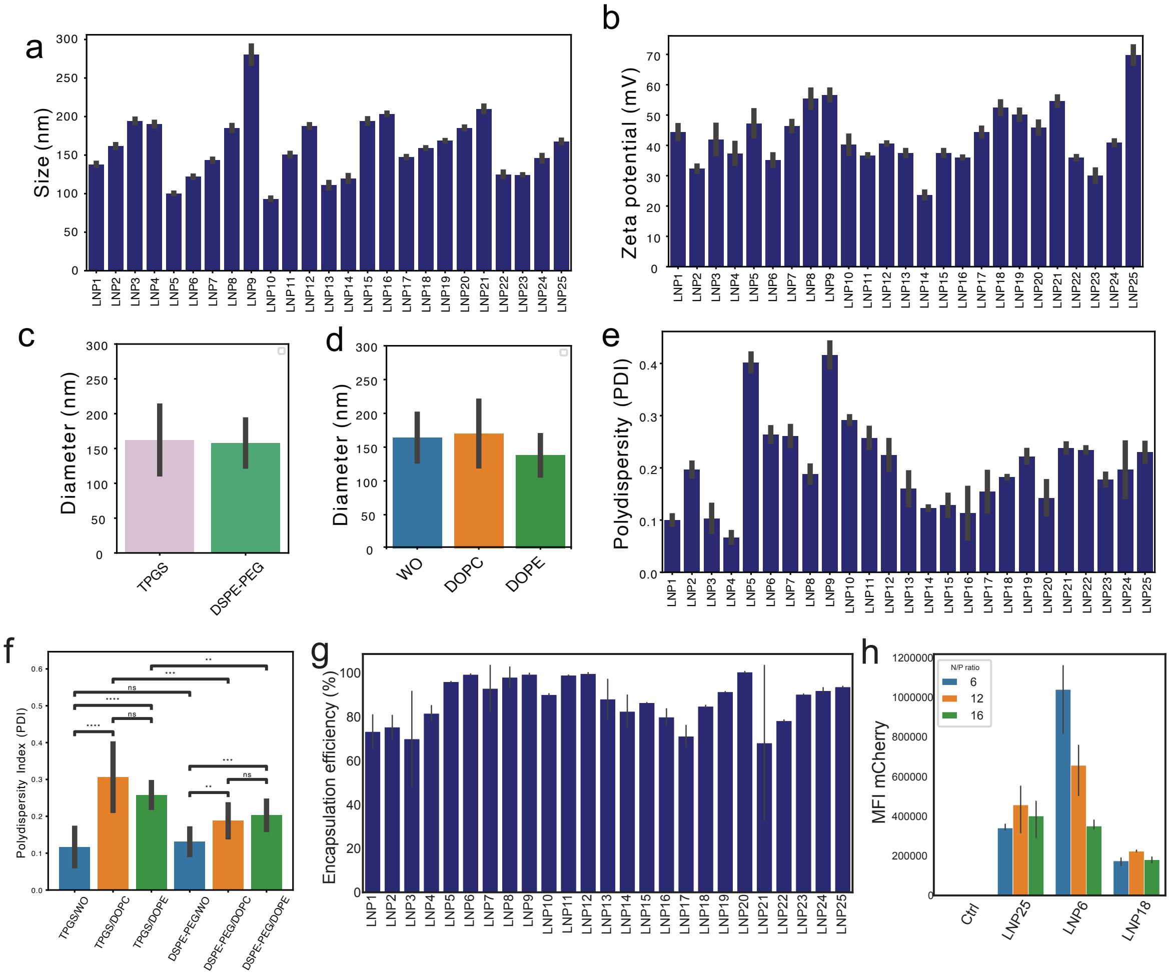


Figure S2. Characterization of LNPs. (a) Average hydrodynamic diameter. (b) Zeta potential of individual LNPs. (c) Average size of LNP library formulated with TPGS versus DSPE-PEG as PEG-lipid. (d) Average size of LNP library formulated with DOPC, DOPE as helper lipid or without helper lipid (WO). (e) Average polydispersity index of individual LNPs. (f) Average polydispersity index of TPGS library without helper lipid (TPGS/WO), DOPC (TPGS/DOPC), DOPE (TPGS/DOPE) and DSPE-PEG2000 particle library without helper lipid (DSPE-PEG/WO), DOPC (DSPE-PEG/DOPC), and DOPE (DSPE-PEG/DOPE). Data are presented as Mean+- SD (n = 12 for TPGS/WO and DSPE-PEG/WO; n = 15 for TPGS/DOPC and DSPE-PEG/DOPC; n = 9 for TPGS/DOPE and DSPE-PEG/DOPE groups); * $p < 0.05$, ** $p < 0.01$, *** $p < 0.001$. Significance was determined by independent t-test. (g) Encapsulation efficiencies of the LNPs for mRNA at N/P ratio of 12. (h) Mean fluorescent intensity (MFI) of HEK293-H cells transfected with selected

LNPs (LNP25, LNP6, LNP18) encapsulated with mRNA (mCherry) at N/P ratio of 6, 12, and 16, respectively.

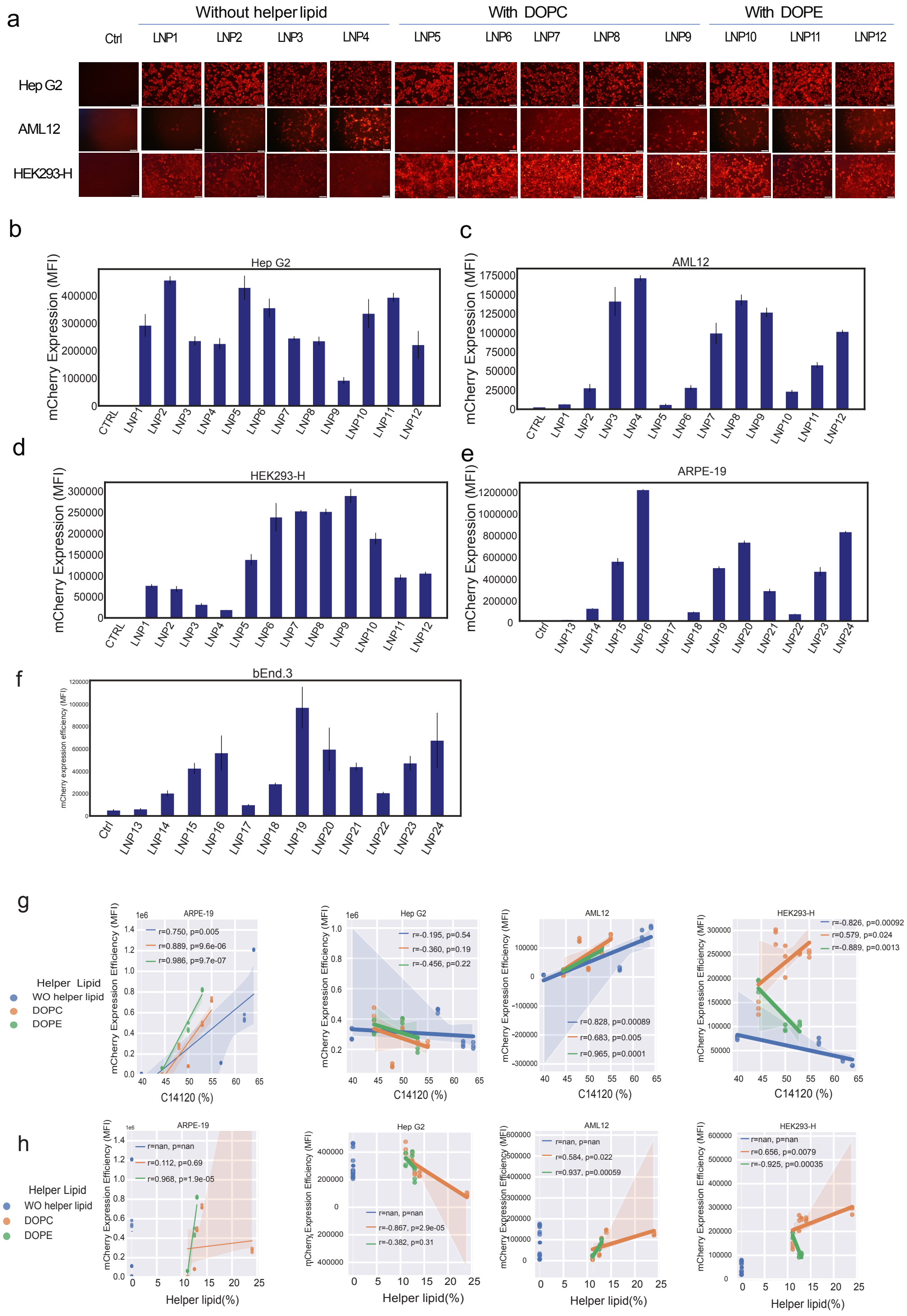

Figure S3. In vitro mRNA expression (mCherry) in various cell types and their correlation with the lipid compositions of LNPs. (a) Representative fluorescent images of mCherry expression in HepG2, AML12 and HEK293H cells transfected with LNP1-LNP12. (b) Quantification of mCherry expression in HepG2, (c) AML12, (d) HEK293H, and (e) ARPE19 by FACS. Data are presented as Mean+- SD (n = 3). Data are recorded with 10000 events for each sample from b to e. (f) Quantification of mCherry expression in bEnd.3 by FACS. Data are presented as Mean+- SD (n = 3). With some samples recorded with events lower than 5000. (g) The mCherry coding mRNA expression level as a function of C14120 percentages and (h) helper lipid % in ARPE19, Hep G2, AML12, and HEK293-H cells. FACs data from bEnd.3 cells was not further analysed for regression fit due to low events acquisition.

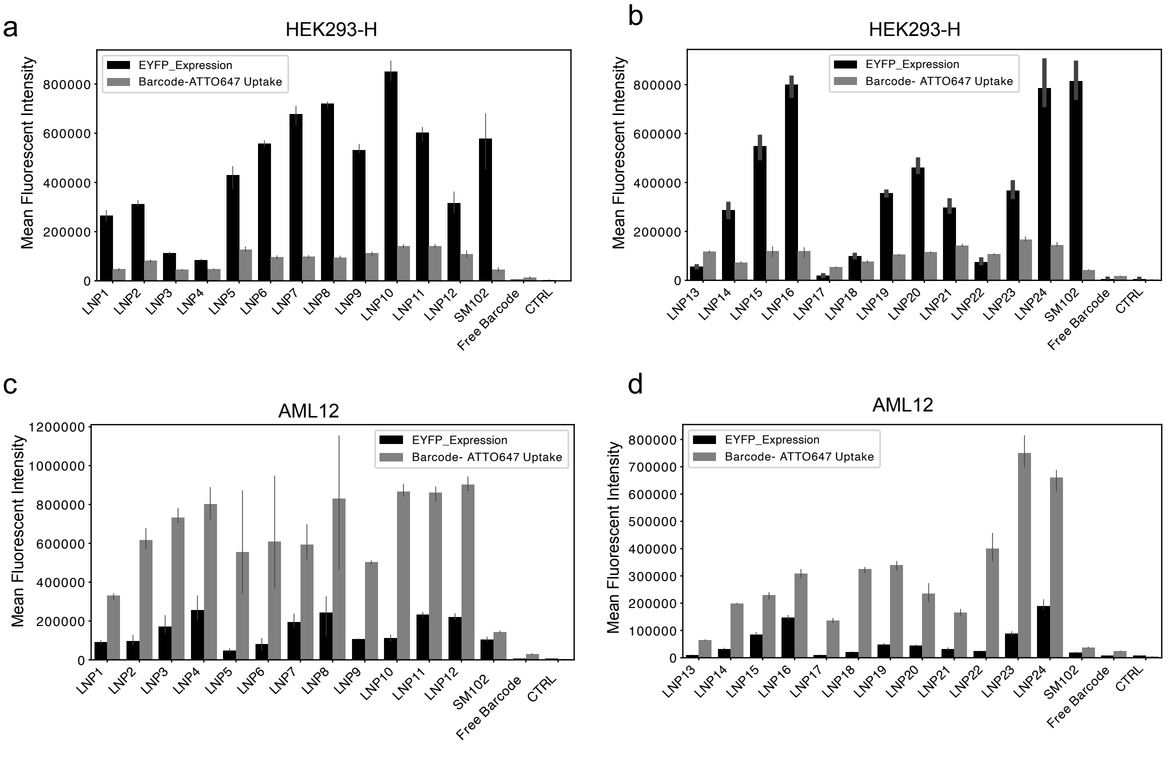


Figure S4. Codelivery of EYFP mRNA and DNA barcode (labelled with ATTO647) in HEK293-H and AML12 cells. EYFP mRNA expression levels (black) versus barcode uptake (grey) in HEK293-H (a-b)

and AML12 (c-d) cells using LNP1-24, compared with SM102 as a positive control and free barcode and untreated cells (CTRL) as negative controls. Data are presented as Mean+- SD (n = 3).

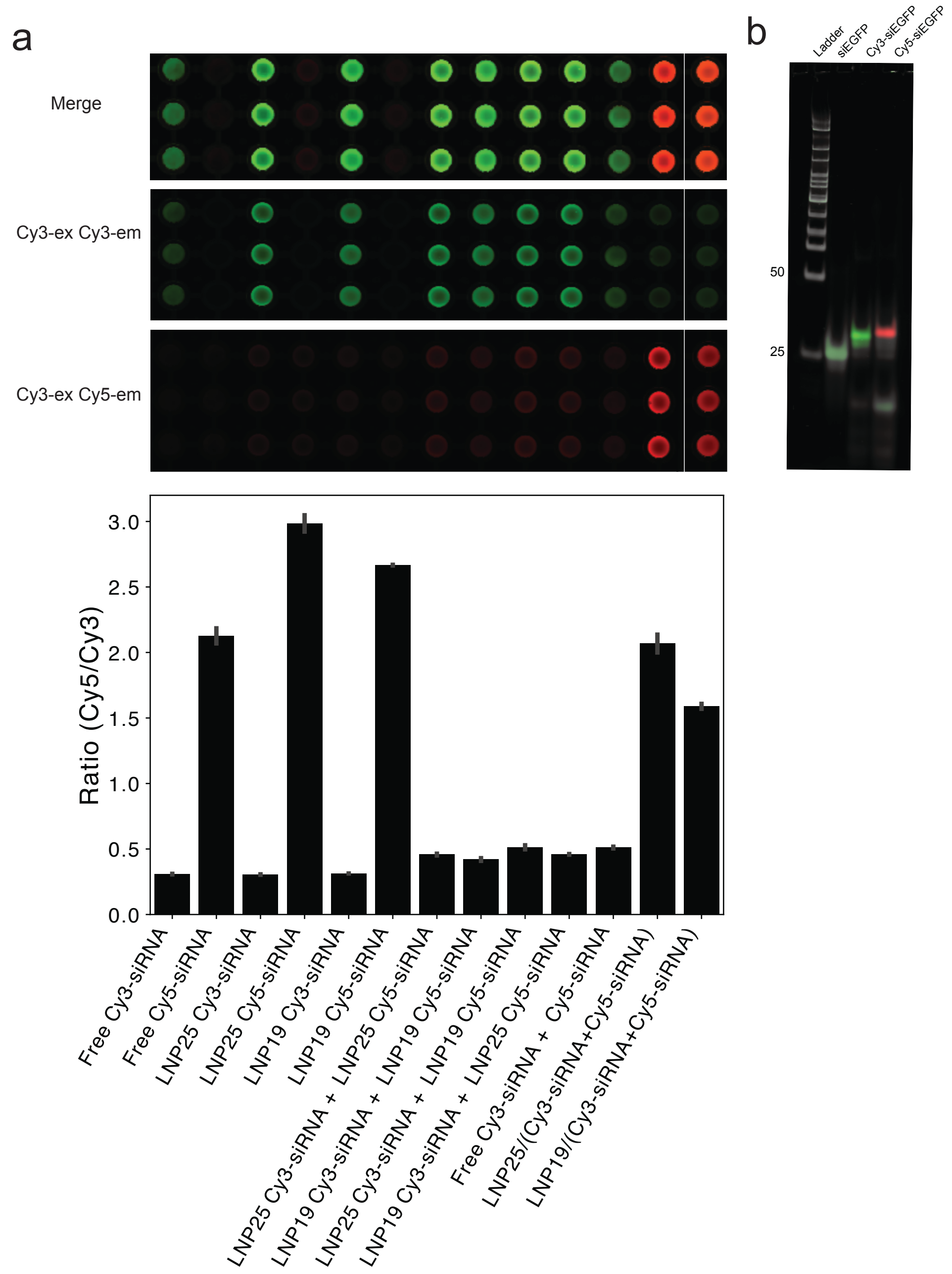


Figure S5. (a) Evaluation of fusion between particles by FRET. LNP19 and LNP25 were randomly selected and encapsulated at N/P ratio of 12 with either Cy3-siRNA or Cy5-siRNA, respectively, or co-encapsulation of both. Upon mixing, merged fluorescent image was recorded by Typhoon scanner by excitation for Cy3 while emission for both Cy3 (Green) and Cy5 (Red). FRET from ratio of Cy5/Cy3 was used as indicative of particle fusion. White line marks the discontinuous of original image. (b) 10% native PAGE gel electrophoresis of siRNA samples labeled either with Cy3 or Cy5.

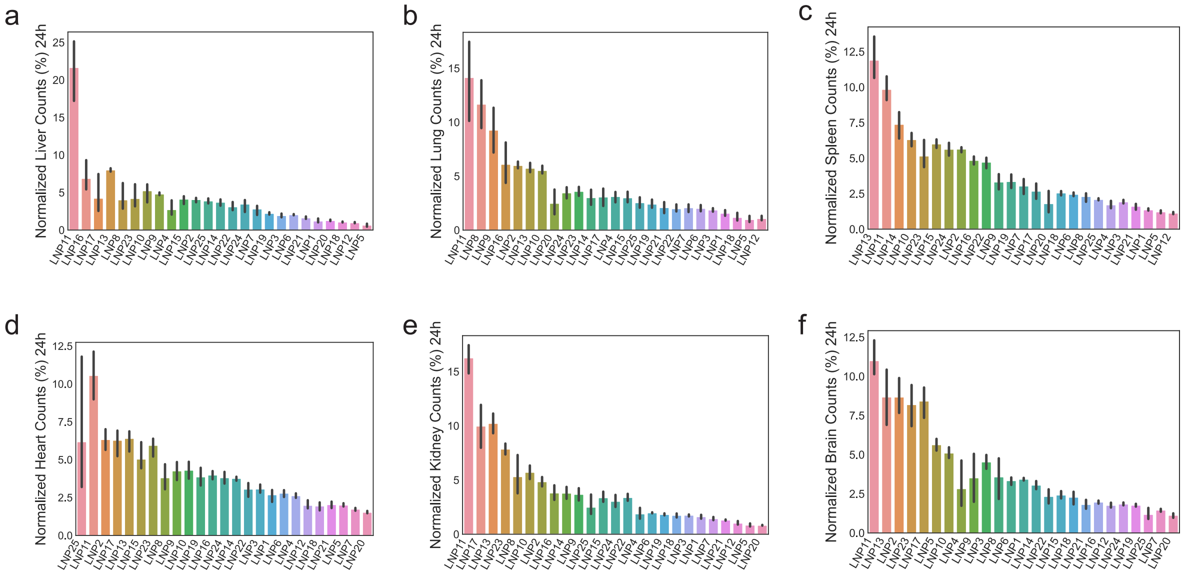


Figure S6. Ranking of LNP accumulation within (a) liver, (b) lung, (c) spleen, (d) heart, (e) kidney, and (f) brain tissues.

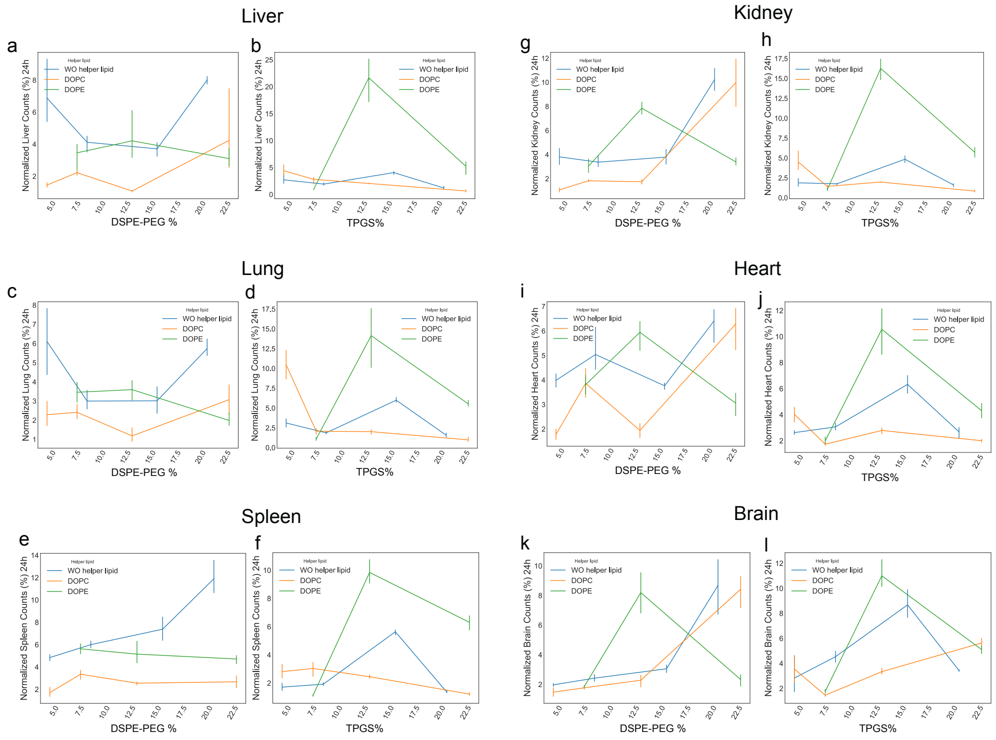


Figure S7. The normalized DNA counts in relation to the lipid percentage present in the formulations in various organs. The normalized DNA counts in relation to DSPE-PEG percentage and TPGS percentage in (a, b) liver; (c,d) lung; (e, f) spleen; (g, h) kidney; (i, j) heart, and (k, l) in brain tissues.

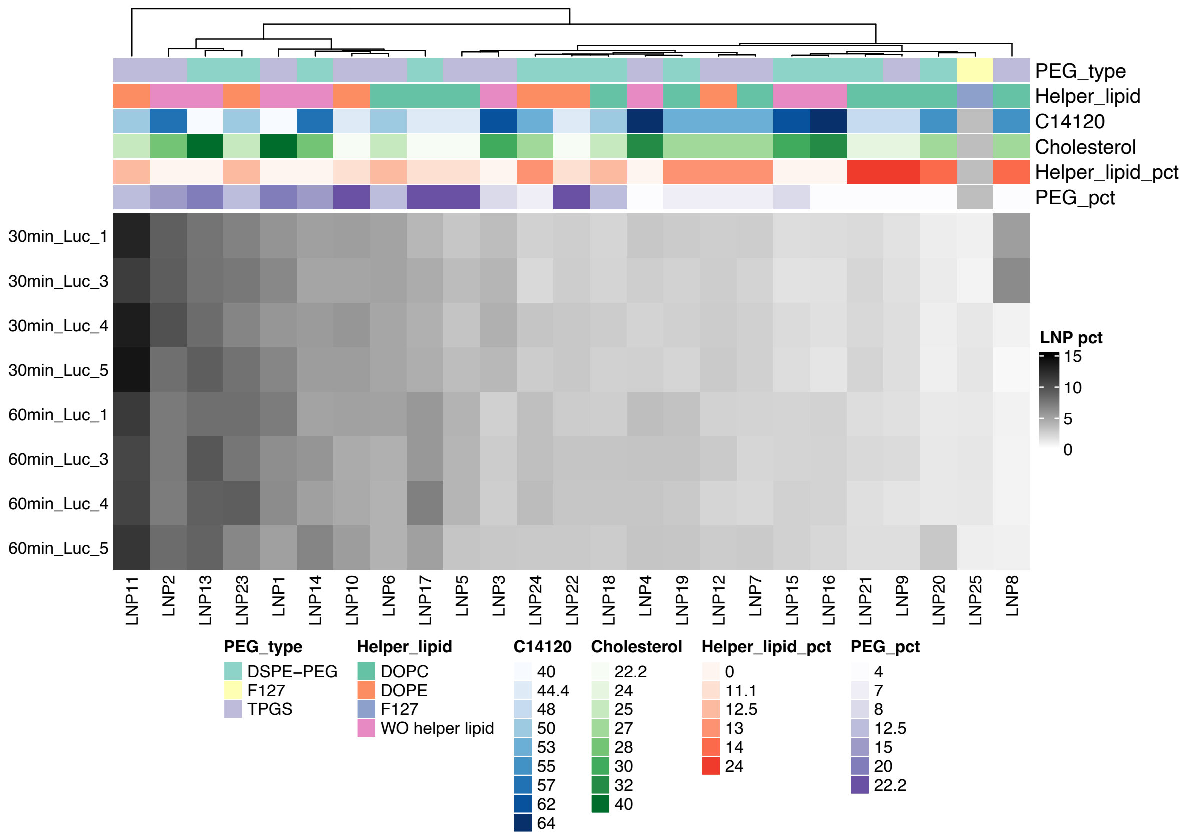


Figure S8. Relative distribution of LNPs in blood circulation over time. The LNPs were quantified by profiling the DNA barcodes by sequencing. Heatmap showing the normalized DNA counts in serum measured at 30 min and 60 min after i.v. injection of LNPs. Note that quantification of absolute differences in LNP levels between time points was not feasible, due to equimolar sequencing for each sample.

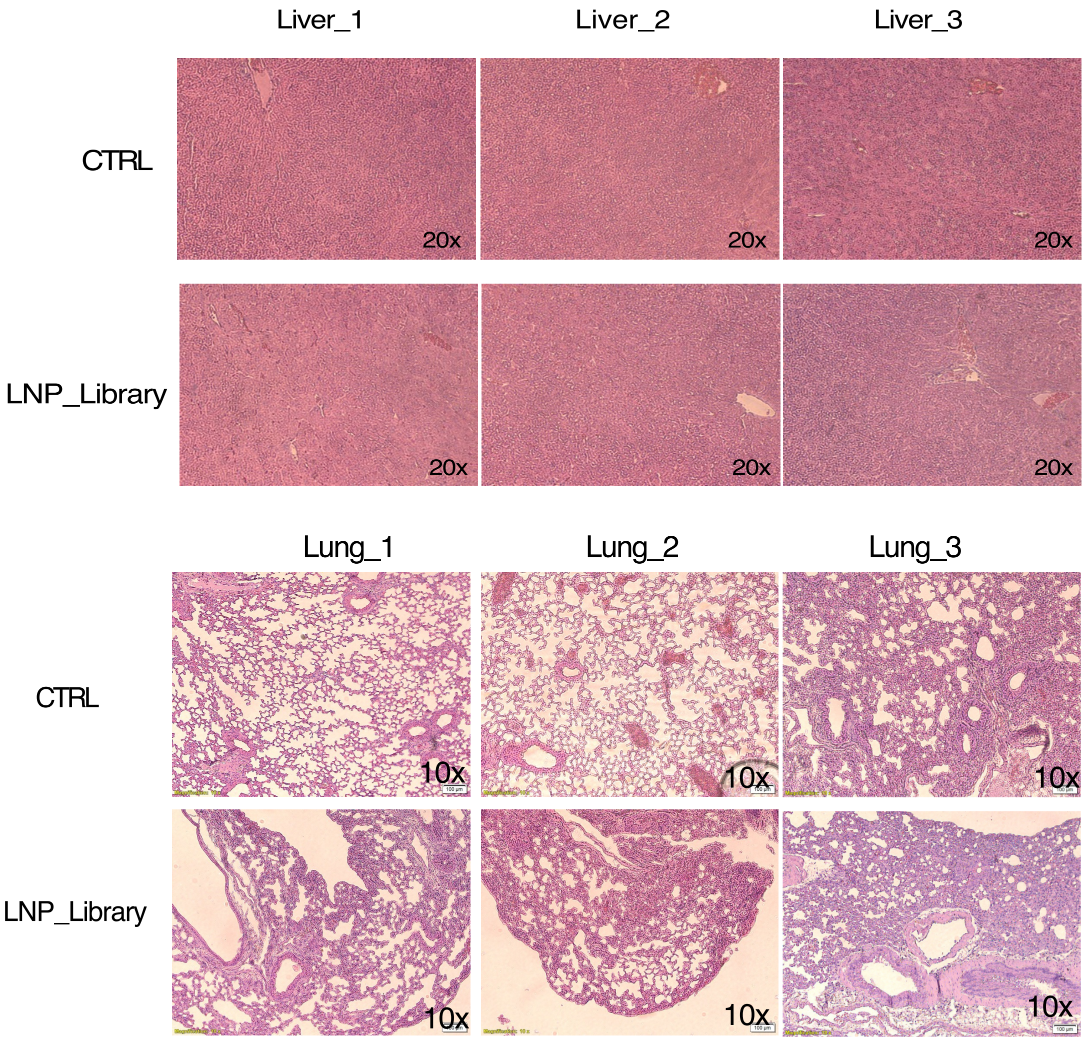


Figure S9. Representative images of H&E staining of liver and lung sections from mice treated with LNP library and control mice treated with PBS (CTRL).

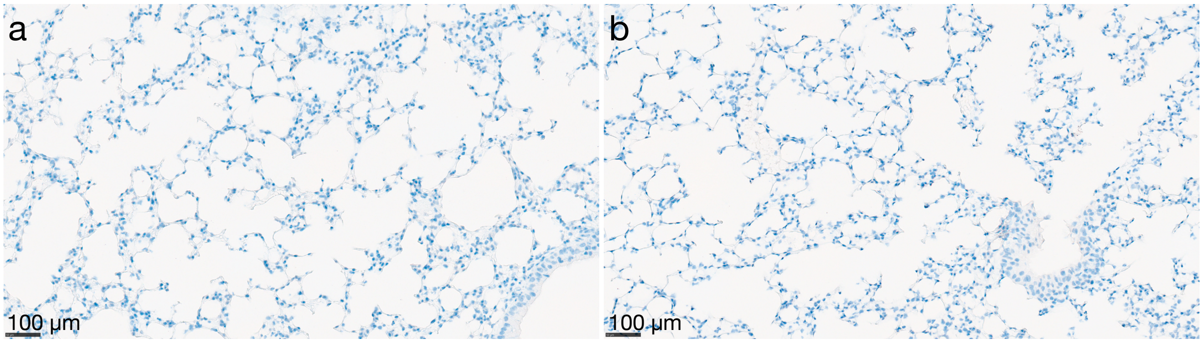


Figure S10. mCherry staining in lung tissue sections from control mice treated with PBS. Representative immunohistochemical staining for mCherry in normal lung tissue sections from two individual control mice (a and b). No mCherry-positive staining was observed. Sections were counterstained with hematoxylin to visualize nuclei. Scale bars = 100 µm (indicated in the lower left corner).

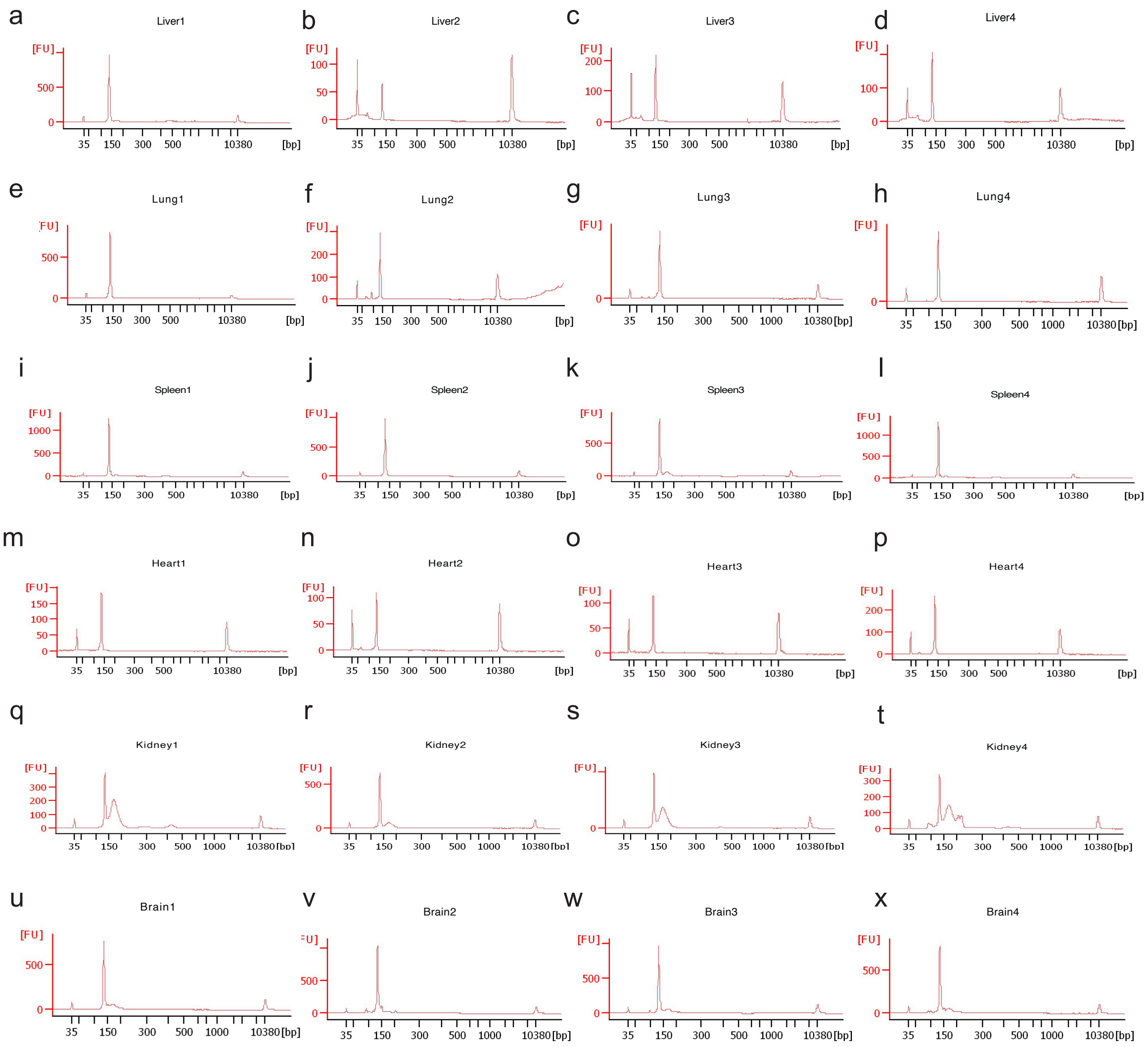


Figure S11. Sample purity assessed by Bioanalyser 2100 prior to pooling for NGS sequencing. DNA barcodes extracted from diverse tissues underwent PCR amplification with adapter sequences and subsequent purification with AMpure beads. The purity of samples was determined using the Bioanlalyser 2100, with representative peak 127nt observed for liver (a-d), lung (e-h), spleen (i-l), heart (m-p), kidney (q-t), and brain (u-x) tissues.
